# Electrophysiological Correlates of Reward Processing in the Human Ventral Tegmental Area

**DOI:** 10.64898/2026.01.27.701696

**Authors:** Arjun Ramaswamy, J. Douglas Steele, Jonathan P. Roiser, Lucy Simmonds, Susie Lagrata, Manjit S. Matharu, Umesh Vivekananda, Harith Akram, Ludvic Zrinzo, Vladimir Litvak

**Author notes:** **Correspondence to:** Prof. Vladimir Litvak, Department of Imaging Neuroscience, UCL Queen Square Institute of Neurology, 12 Queen Square, London, WC1N 3AR, UK.

## Abstract

The ventral tegmental area (VTA) is the principal source of dopaminergic input to the human prefrontal cortex and is hypothesised to play a central role in reward–based learning. However, direct electrophysiological evidence in humans has been lacking. Here, using intracranial recordings obtained during an instrumental learning task in patients undergoing deep brain stimulation surgery for chronic cluster headache (n=13), we show that VTA local field potentials selectively encode rewarding outcomes compared with losses or neutral outcomes independently of whether behavioural learning occurred. In a subset of participants (n= 8) who learned to prefer a high–reward option, VTA responses to unexpected rewards were larger than to expected rewards. In two participants who sufficiently explored both options, VTA activity reflected the expected value of the chosen option during decision–making. These findings provide direct electrophysiological evidence that the human VTA encodes reward–related signals consistent with reinforcement learning mechanisms described in animal models.

---

The ventral tegmental area (VTA) is a key node in the reward and motivation system of the brain. Dopaminergic signalling within this region enables learning from outcomes, behavioural adaptation, and the pursuit of rewarding goals. Disruption of this system contributes to a range of psychiatric and neurological disorders, including addiction, depression, and chronic pain, underscoring the clinical and societal importance of understanding how the human VTA encodes reward and motivational value ^1–3^. Yet despite its central role, direct human electrophysiological evidence for VTA reward signalling remains scarce.

In animal models, seminal work by Schultz and colleagues showed that dopaminergic neurons in the VTA signal reward prediction errors (RPEs) i.e. the difference between expected and actual outcomes serving as teaching signals in temporal-difference learning frameworks ^4,5^. These neurons increase firing when an unexpected reward is delivered and pause when an expected reward is omitted, encoding not simply reward presence but the mismatch between expectation and outcome. Early studies also suggested that VTA dopamine neurons primarily represent reward value rather than loss or aversion, distinguishing reward from punishment encoding ^5^. Subsequent experiments demonstrated that dopamine neuron responses obey the formal principles of associative learning. In particular, phasic activity diminishes when a reward is fully predicted and re-emerges for unpredicted rewards that signal a prediction error ^6^. Moreover, dopamine responses follow the “blocking” rule of the Rescorla–Wagner model, showing no activation to redundant cues that add no new predictive value ^7^. Later work revealed greater heterogeneity in dopaminergic responses ^8^, but the core principle that dopamine neurons convey RPE remains central to reinforcement learning (RL) theory. Animal studies also provide evidence that reward processing is reflected in VTA Local Field Potentials (LFP). Pasquereau et al.^9^ demonstrated that midbrain LFPs reflect reward-correlated information in low-frequency bands, synchronised across neuronal populations with dopaminergic involvement. Oscillatory activity in the VTA also adapts to changing cue-reward contingencies ^10^, and cross-regional synchrony suggests a coordinated role in learning.

In humans, direct electrophysiological evidence is rare, partly due to the VTA’s deep midbrain location and limited clinical access. Indirect approaches have been more informative. fMRI studies show that BOLD activity in the VTA scales with RPE magnitude ^11,12^, while EEG studies link mid-frontal theta oscillations and the feedback-related negativity to dopaminergic error processing ^13^. Intraoperative recordings from the neighbouring substantia nigra have provided the first direct evidence that single neurons respond more strongly to unexpected rewards than to losses ^14^.

Recent causal evidence further highlights the relevance of the VTA to human decision-making. In subjects with trigeminal autonomic cephalalgias undergoing deep brain stimulation (DBS) targeting the VTA, Hirschbichler et al.^15^ found that stimulation increased strategic betting behaviour without impairing reinforcement learning, suggesting that VTA neuromodulation can bias motivational processes.

Complementing these subcortical findings, recent human intracranial studies have begun to map reinforcement-learning computations across cortico-thalamic circuits with high temporal precision. Intracranial high-frequency activity has been shown to encode outcome valence in an expectation-dependent manner across widespread cortical regions during probabilistic learning, consistent with distributed learning-related surprise signals ^16^. Furthermore, high-frequency activity in the ventromedial prefrontal and orbitofrontal cortices has been shown to encode subjective value ^17^. Depth recordings demonstrate prediction-error-related activity in frontal and insular cortex, including dissociable reward- and punishment-linked signals in vmPFC/lateral orbitofrontal versus anterior insula/dorsolateral prefrontal regions^18^, as well as asymmetric coding of positive, negative, and unsigned reward prediction errors in insula and dorsomedial prefrontal cortex^19^. Extending beyond cortex, human thalamic recordings show low-frequency oscillations that correlate with model-derived expected value and outcome during reinforcement learning^20^.

Together, these results motivate direct investigation of the human VTA’s electrophysiological role in reward processing. Here, we used a rare clinical opportunity to record VTA LFPs directly in patients undergoing DBS implantation for severe, drug-resistant chronic cluster headache, a subtype of trigeminal autonomic cephalalgia. During temporary electrode externalisation, participants performed a probabilistic instrumental learning task while we recorded VTA LFPs. This design allowed us to test whether population-level VTA activity in humans reflects reward-learning dynamics predicted by reinforcement learning theory.

We tested two related but distinct hypotheses

i. At the cohort level, we hypothesised that VTA local field potentials would differentiate rewarding outcomes from non-rewarding outcomes, but would not differentiate loss outcomes from neutral outcomes, consistent with reward-selective rather than general salience encoding ^2,3^. Outcome-related analyses were conducted across all participants and did not assume successful behavioural learning.
ii. In participants demonstrating clear behavioural evidence of reward learning, we hypothesised that VTA activity would correlate with model-derived expected value estimates. Model-based analyses were therefore restricted to participants meeting predefined behavioural learning criteria.

## Results

Fourteen subjects (9 male, age 46±10) undergoing DBS surgery for chronic cluster headache treatment were recruited (see Supplementary Table S1 for details). Figure 1A shows the study flow chart and Figure 1B shows electrode locations in relation to a template structural image. The subjects were asked to perform an instrumental learning task (Figure 2A) while their VTA-LFPs were recorded. On each trial, subjects selected one of two fractal stimuli, each probabilistically associated with different outcomes. There were three different trial types, each associated with a fixed fractal pair, randomised across subjects. In reward trials, one fractal had a 0.7 probability of leading to a win (“You win”) and a 0.3 probability of no outcome (“Nothing”), while the other fractal had the opposite contingency (0.3 probability of winning, 0.7 probability of nothing). Loss trials followed the same structure but with aversive reinforcement, where one fractal had a 0.7 probability of loss (“You lost”) and a 0.3 probability of no outcome. Neutral trials provided no reinforcement, with equal probabilities (0.5/0.5) of “Look” or “Nothing.” Subjects were instructed to maximize reward and minimize loss through trial and error, as outcome probabilities remained fixed but were unknown to them at the start of the task.

**Figure 1.**
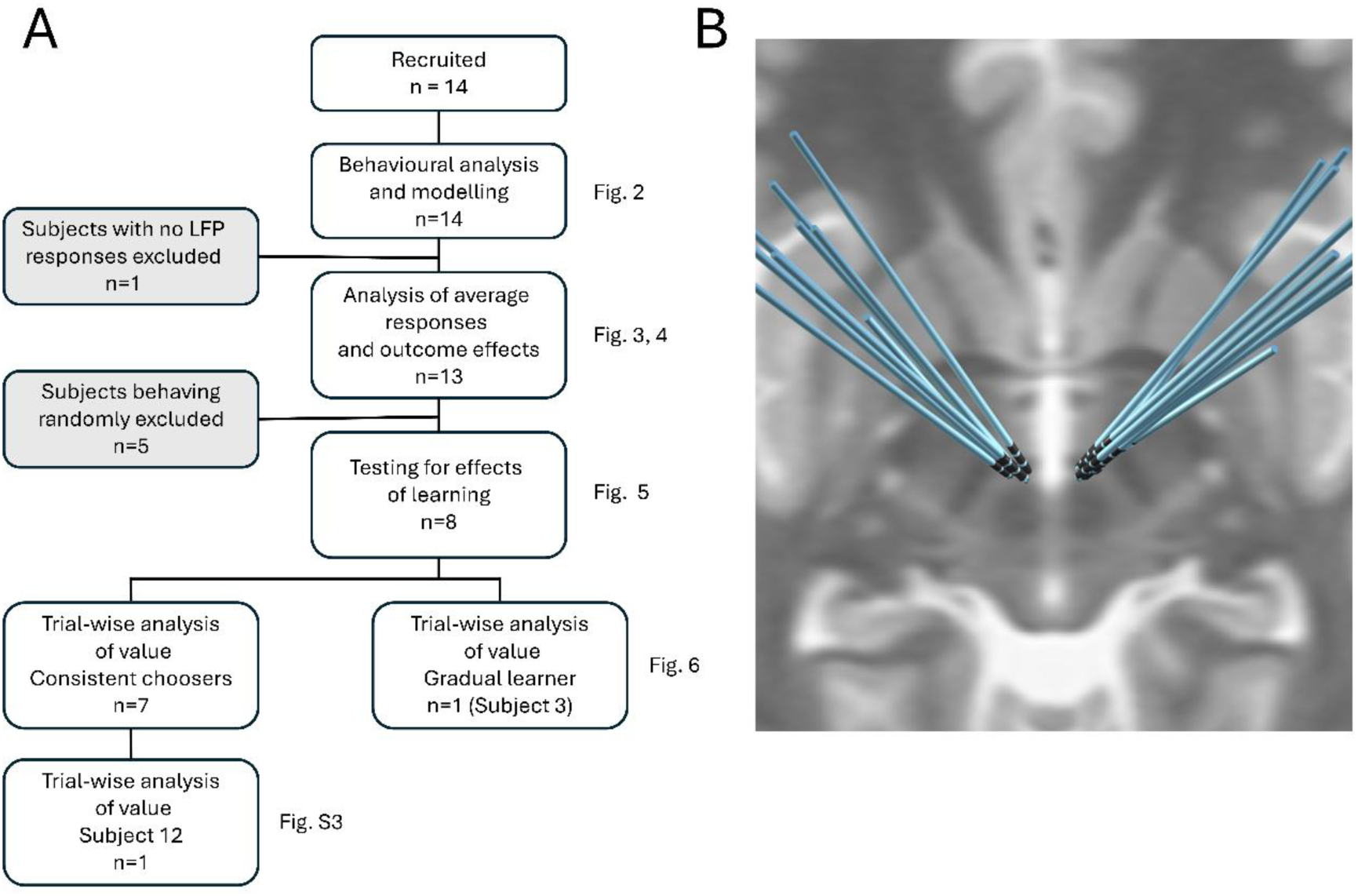
(A) Flow diagram for the study. See the main text for details. (B) VTA electrode locations reconstructed with Lead-DBS overlaid on a T2 template image.

**Figure 2.**
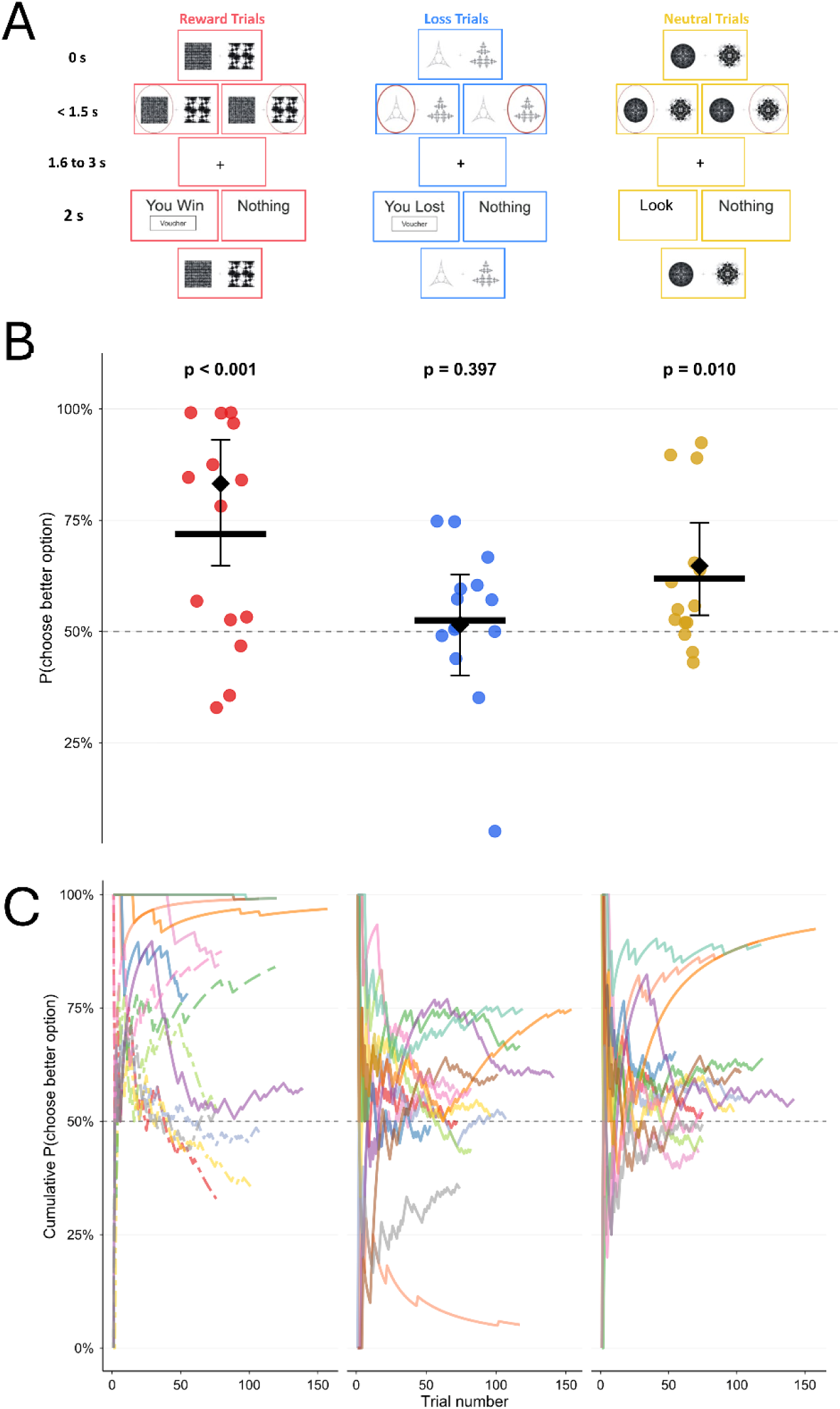
(**A**) **Task paradigm for the instrumental learning task.** On each trial, subjects selected one of two fractal stimuli, each probabilistically associated with different outcomes. In reward trials, one fractal had a 0.7 probability of leading to a win (“You win”) and a 0.3 probability of no outcome (“Nothing”), while the other fractal had the opposite contingency (0.3 probability of winning, 0.7 probability of nothing). Loss trials followed the same structure but with aversive reinforcement, where one fractal had a 0.7 probability of loss (“You lost”) and a 0.3 probability of no outcome. Neutral trials provided no reinforcement, with equal probabilities (0.5/0.5) of “Look” or “Nothing.” Each trial began with the presentation of a fractal pair for 1.5 seconds, during which subjects had to make a response. If no response was made within this 1.5-second window, the trial was skipped. Following a response, there was a variable delay of 1.6 to 3 seconds before outcome feedback was presented. The outcome remained on-screen for 2 seconds. An inter-trial interval (ITI) of 2 seconds separated consecutive trials. For Subject 1, the interval between button presses and outcome presentation ranged from 4 to 6 seconds, while for Subject 2, it ranged from 3.5 to 6 seconds. After these initial subjects, this interval was standardized to 1.6 to 3 seconds. Additionally, the ITI was initially jittered in the first two subjects but was later fixed at 2 seconds to save time. Subjects were instructed to maximize reward and minimize loss through trial and error, as outcome probabilities remained fixed but were unknown to them at the start of the task. **B. Model-free behavioural validation.** Each dot represents one subject’s proportion of choosing the better option (or option 1 for neutral). The crossbar indicates the group mean; the black diamond shows the GLMM-estimated group probability with 95% confidence intervals. The dashed horizontal line marks chance performance (50%). Reward trials showed significantly above-chance performance (one-sided p < 0.001), loss trials did not (one-sided p = 0.397), and neutral trials showed a significant systematic preference (two-sided p = 0.010). **C. Cumulative learning curves for all 14 subjects across the three trial types.** Each line represents one subject, colour-coded by subject identity. In the reward panel, line type distinguishes behavioural type assigned from model comparison: solid = Consistent (C), long-dash = Gradual (G), dotted = Random (R). The dashed horizontal line indicates chance (50%)

Below we will use ‘Reward’, ‘Loss’ and ‘Neutral’ to denote the trial types and ‘win’, ‘loss’, ‘look’ and ‘nothing’ to denote the possible outcome types.

### Trial completion and model-free performance

All subjects completed the task, contributing a total of 4,183 valid trials (1,414 reward, 1,385 loss, 1,384 neutral). Trial counts per subject ranged from 163 to 469 (mean ± SD: 298.8 ± 86.9; Table 1). Mixed effects logistic regression with random subject intercepts confirmed that group level choice accuracy was significantly above chance in reward trials (estimated P = 83.2%, 95% CI [64.8, 93.0], z = 3.17, one-sided p = 0.0008), but not in loss trials (estimated P = 51.5%, 95% CI [40.1, 62.8], z = 0.26, one-sided p = 0.397). A two-sided test for neutral trials revealed a significant systematic preference (estimated P = 64.7%, 95% CI [53.6, 74.4], z = 2.58, p = 0.01), though this likely reflects stimulus or side bias rather than learning given the 50/50 contingency. Individual binomial tests showed that 8/14 subjects performed better than chance for Reward trials, 5/14 for Loss and 6/14 for Neutral. These results are summarised in Figure 2B and Table 1.

**Table 1.** Model-agnostic behavioural results and trial numbers.

| Subject ID | Choice rates (%) and individual binomial tests |  |  |  |  |  | Trial numbers |  |  |  |
| --- | --- | --- | --- | --- | --- | --- | --- | --- | --- | --- |
|  | High reward chosen | p | Low loss avoided | p | Neutral option 1 | p | Rew | Loss | Neu | Total |
| 1 | 32.9 | 0.999 | 50.0 | 0.547 | 52.0 | 0.818 | 76 | 72 | 75 | 223 |
| 2 | 78.2 | **<0.001\*** | 49.1 | 0.608 | 65.5 | **0.030\*** | 55 | 53 | 55 | 163 |
| 3 | 84.0 | **<0.001\*** | 66.7 | **<0.001\*** | 63.9 | **0.003\*** | 119 | 117 | 119 | 355 |
| 4 | 56.8 | 0.063 | 59.6 | **0.014\*** | 54.9 | 0.275 | 139 | 141 | 142 | 422 |
| 5 | 96.8 | **<0.001\*** | 74.7 | **<0.001\*** | 92.4 | **<0.001\*** | 157 | 154 | 158 | 469 |
| 6 | 99.1 | **<0.001\*** | 60.4 | **0.023\*** | 61.2 | **0.030\*** | 107 | 101 | 103 | 311 |
| 7 | 87.5 | **<0.001\*** | 57.3 | 0.112 | 52.7 | 0.728 | 80 | 82 | 74 | 236 |
| 8 | 52.6 | 0.366 | 35.1 | 0.997 | 49.3 | 1.000 | 76 | 74 | 75 | 225 |
| 9 | 99.2 | **<0.001\*** | 74.8 | **<0.001\*** | 89.0 | **<0.001\*** | 120 | 119 | 118 | 357 |
| 10 | 99.2 | **<0.001\*** | 5.1<sup>†</sup> | 1.000 | 89.7 | **<0.001\*** | 120 | 117 | 116 | 353 |
| 11 | 46.8 | 0.778 | 50.5 | 0.500 | 55.8 | 0.281 | 109 | 107 | 104 | 320 |
| 12 | 84.6 | **<0.001\*** | 57.1 | 0.141 | 43.1 | 0.289 | 78 | 70 | 72 | 220 |
| 13 | 53.3 | 0.324 | 43.9 | 0.888 | 45.3 | 0.489 | 77 | 82 | 75 | 234 |
| 14 | 35.6 | 0.999 | 51.0 | 0.459 | 52.0 | 0.762 | 101 | 96 | 98 | 295 |
| **Total** |  |  |  |  |  |  | **1414** | **1385** | **1384** | **4183** |
| **Sig.** |  | **8/14** |  | **5/14** |  | **6/14** |  |  |  |  |
| **Mean±SD** |  |  |  |  |  |  | 101.0±28.6 | 98.9±28.8 | 98.9±29.7 | 298.8±86.9 |

Choice rates indicate the percentage of trials on which the high-reward option was chosen (reward), the high-loss option was avoided (loss), or option 1 was selected (neutral). Individual p-values are from exact binomial tests against chance (50%): one-sided (greater than chance) for reward and loss, two-sided for neutral. Significant results ( $p < 0.05$ ) are marked with \* and shown in bold. Trial counts are shown per subject and condition (reward: 70/30 win contingency; loss: 70/30 loss contingency; neutral: 50/50). <sup>†</sup>Subject 10 consistently chose the high-loss option in loss trials.

### Reaction times

Reaction times (RTs) were analysed to assess task engagement. A repeated-measures ANOVA revealed a significant effect of trial type (Reward/Neutral/Loss) on RTs (F(2,26) = 7.49, p = 0.0027). Post-hoc comparisons (Bonferroni corrected) showed that subjects responded significantly faster during Reward trials (M = 0.83 s, SD = 0.32 s) compared to Loss trials (M = 0.92 s, SD = 0.331 s; p = 0.0049). No significant differences were observed between Reward and Neutral trials (p = 0.34) or Loss and Neutral trials (p = 0.086).

### Model comparison for reward trials

To further characterise individual differences in learning strategy, we compared three models: random choice (Null), Rescorla-Wagner with a single learning rate (RW-1α), and Rescorla-Wagner with dual learning rate (RW-2α), using per-subject Bayesian Information Criterion (BIC, Table 2). The Null model provided the best fit for five subjects (1, 8, 11, 13, 14), indicating random or disengaged responding. These were all subjects for whom individual binomial tests indicated random behaviour in the model-free analysis. The single-learning-rate model (RW-1α) was preferred for two subjects (3 and 7), both of whom displayed gradual, exploratory learning: these subjects initially sampled both options and progressively converged on the high-value choice. For Subject 3, the fitted learning rate was α = 0.065 with β = 7.03, indicating slow but stable value accumulation with highly deterministic choice. The dual-learning-rate model (RW-2α) was preferred for seven subjects (2, 4, 5, 6, 9, 10, 12), who showed consistent, near-ceiling choice of the high-reward option (Subject 4 was borderline in the binomial test with p = 0.063). These subjects were characterised by high reward learning rates (α_rew_ = 0.33–0.80), very low non-rewarding outcome learning rates (α_nothing_ = 0.01–0.04), and high inverse temperatures (β = 4.07–9.38), reflecting strong asymmetric learning and near-deterministic choice.

**Table 2.** Model comparison results for reward trials.

| Subject | Behavioural type | BIC Null | BIC RW-1α | BIC RW-2α | Best model | $\alpha_{\text{rew}}$ | $\alpha_{\text{nothing}}$ | $\beta$ |
| --- | --- | --- | --- | --- | --- | --- | --- | --- |
| 1 | R | **105.4** | 114.5 | 118.6 | Null | — | — | — |
| 2 | C | 76.2 | 72.1 | **67.5** | RW-2α | 0.507 | 0.038 | 4.07 |
| 3 | G | 165.0 | **110.4** | 114.3 | RW-1α | 0.065* | — | 7.03 |
| 4 | C | 192.7 | 200.9 | **185.8** | RW-2α | 0.746 | 0.011 | 4.19 |
| 5 | C | 217.6 | 62.1 | **61.6** | RW-2α | 0.607 | 0.025 | 7.17 |
| 6 | C | 148.3 | 53.9 | **25.5** | RW-2α | 0.328 | 0.014 | 9.38 |
| 7 | G | 110.9 | **68.4** | 68.7 | RW-1α | 0.080* | — | 9.08 |
| 8 | R | **105.4** | 114.1 | 118.2 | Null | — | — | — |
| 9 | C | 166.4 | 54.8 | **26.7** | RW-2α | 0.663 | 0.022 | 9.25 |
| 10 | C | 166.4 | 42.7 | **26.5** | RW-2α | 0.580 | 0.020 | 9.25 |
| 11 | R | **151.1** | 160.8 | 165.4 | Null | — | — | — |
| 12 | C | 108.1 | 81.8 | **60.3** | RW-2α | 0.798 | 0.033 | 8.16 |
| 13 | R | **106.7** | 115.7 | 117.2 | Null | — | — | — |
| 14 | R | **140.0** | 149.7 | 154.1 | Null | — | — | — |
| Total best for |  | **5** | **2** | **7** |  |  |  |  |

Bayesian Information Criterion (BIC) values are shown for three models fitted to each subject's reward trial data: a Null model assuming random responding (0 parameters), a single-learning-rate Rescorla–Wagner model (RW-1α ; 2 parameters: $\alpha$ , $\beta$ ), and a dual-learning-rate Rescorla–Wagner model (RW-2α ; 3 parameters: $\alpha_{\text{rew}}$ , $\alpha_{\text{nothing}}$ , $\beta$ ). The lowest (best) BIC for each subject are shown in bold. Based on the winning model and individual choice trajectories, each subject was assigned a behavioural type: R = Random (Null wins), indicating no evidence of contingency-sensitive responding; G = Gradual (RW-1α wins), indicating exploratory learning with progressive convergence on the high-value option; C = Consistent (RW-2α wins), indicating near-ceiling choice of the high-reward option with minimal exploration. Fitted parameters are shown for subjects where RL models were preferred; for RW-1α subjects, a single $\alpha$ was fitted to all outcomes. The RW-2α model was preferred for the majority of engaged subjects (7/14), while 2/4 gradual learners were best described by the simpler RW-1α model and the rest of the subjects (5/14) were best described by the Null model.

The asymmetry between α_rew_ and α_nothing_ (ratios ranging from 13 to 68) indicates that these subjects updated their value estimates strongly after wins but barely after ‘nothing’ outcomes, effectively implementing a win-stay strategy captured within the RL framework. Figure 2C shows cumulative learning curves for all subjects across the three trial types, with reward-trial trajectories differentiated by behavioural type. The individual modelling results are provided in Table 2 and individual choice trajectories, and model fits are shown in Supplementary Figure S1.

### Mean LFP responses

We first examined the mean VTA-LFP responses to each of the three events in the trial: stimulus presentation, decision (button press) and outcome (feedback) presentation. The responses were averaged over trial types and outcome labels. Visual inspection of the individual evoked responses showed that subject 7 had no clear responses to any of the task events. They were, therefore, excluded from further analyses. The results for the remaining 13 subjects are shown in Figure 3. Clear evoked responses appeared during stimulus presentation (significant clusters 0.167–0.317 s, 0.667–0.8 s) and outcome presentation (0.15–0.3 s), but no consistent phase-locked response to button presses was detected at the group level. Significant clusters were identified using family-wise error correction at the cluster level (p<0.05), a standard approach for controlling false positives across multiple time points without requiring assumptions about the timing of effects. We also examined induced responses (changes in oscillatory power not phase-locked to the stimulus) around the same events, but these primarily reflected the time-frequency representation of the evoked responses, as evidenced by close alignment of significant power modulations and inter-trial coherence increases, or occurred later in time (Supplementary Figure S2). We therefore focused exclusively on evoked responses, which represent the phase-locked voltage deflections time-locked to each task event, in our subsequent analysis.

**Figure 3.**
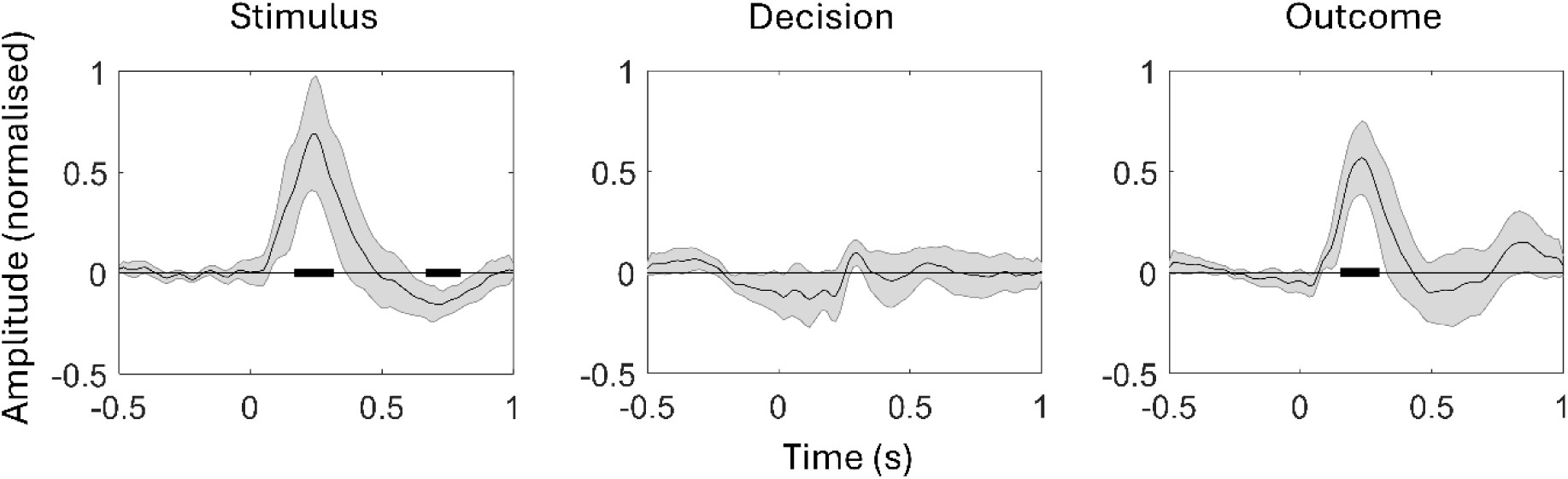
Average VTA evoked responses to the task events. The shaded error bars denote 95% confidence interval for the mean. Black bars denote clusters significantly different from zero (FWE-corrected at the cluster level, p<0.05).

### Analysis of the response to outcome

We next examined outcome-related evoked responses, analysing the three trial types separately and comparing mean responses as well as the difference between different outcomes.

To increase sensitivity, we first identified time windows of interest pooling responses across trial types and then statistically interrogated LFP amplitudes averaged over these windows using a linear mixed effects model (LMM). Note that this does not constitute double dipping^21^ but is akin to the use of orthogonal contrast^22^.

Testing for the difference between ‘win/loss/look’ and ‘nothing’ pooled across trial types, revealed a significant cluster at 0.317-0.467 s post outcome presentation (Figure 4A).

**Figure 4.**
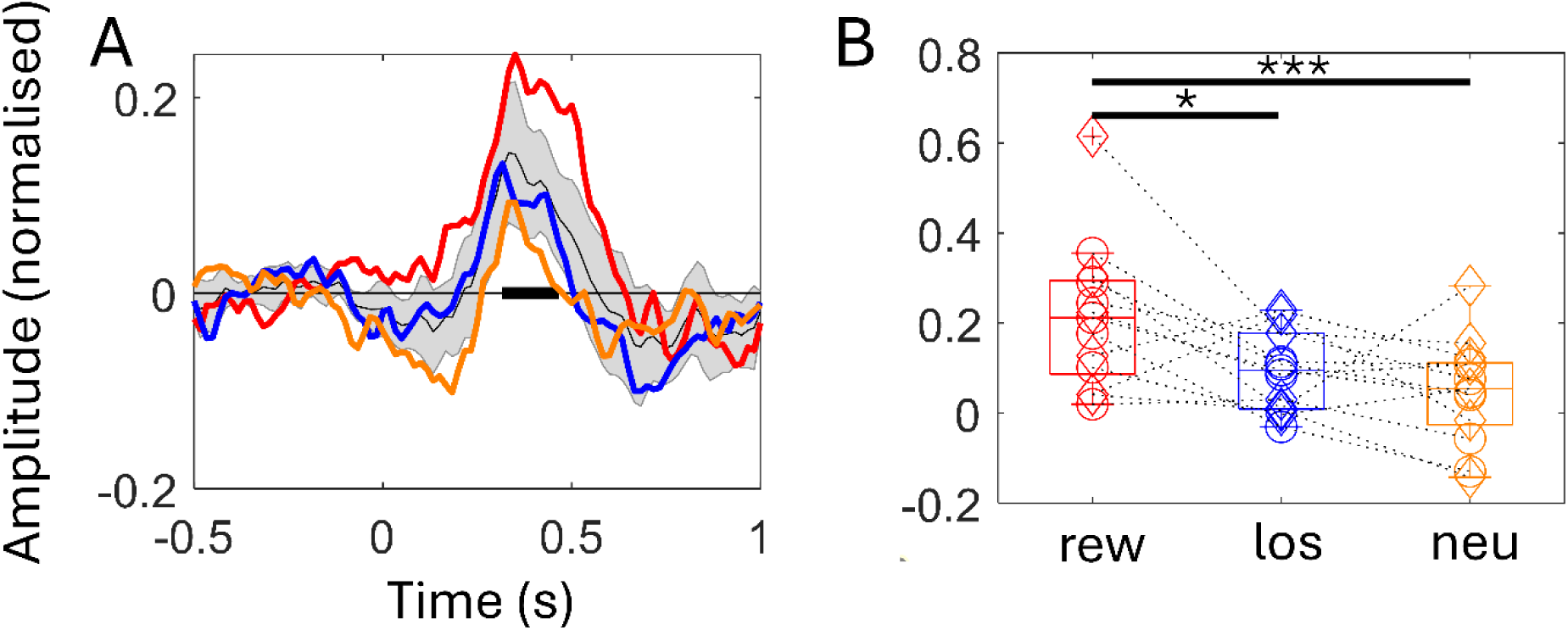
VTA responses to outcome. (**A**) Difference waveform between ‘win’/’loss’/’look’ and ‘nothing’ outcome. The thin black line shows the average response pooled across trial type with the shaded error bars around it representing 95% confidence interval for the mean The black horizontal bar indicates a cluster significantly different from zero (0.317-0.467 s, FWE-corrected at the cluster level, p<0.05). The coloured lines represent separate responses for three trial types: Reward (red), Loss (blue) and Neutral (orange). (**B**) Difference waveforms shown in (A) averaged over the significant cluster in (A) plotted separately for the subjects and trial types. Marker type indicates the behavioural type based on model-agnostic analysis (cf. Table 1): random (diamond) or non-random (circle). On each box, the central mark is the median, the edges of the box are the 25th and 75th percentiles, the whiskers extend to the most extreme datapoints the algorithm considers to be not outliers, and the outliers are plotted individually (see Matlab’s boxplot function). The dotted lines connect points pertaining to the same subject. The black horizontal bars indicate significant differences (Linear Mixed Model, * - p < 0.05, *** - p ≤ 0.001).

When LMM was applied to LFP amplitudes averaged over this window, model comparison (BIC) favoured a parsimonious specification with Trial_Type × Outcome_Label as the core fixed effects. All three trial types showed a positive effect of outcome label: win-nothing (β = 0.342, SE = 0.041, df = 3838, t = 8.33, p < 1×10⁻⁴), loss-nothing (β = 0.158, SE = 0.041, df = 3837, t = 3.86, p = 0.0003), look-nothing (β = 0.118, SE = 0.042, df = 3837, t = 2.85, p = 0.013).

A test of the null hypothesis that the outcome effects were equal for the three trial types was significant (F(2, 3836.2) = 8.46, p = 0.0002). This indicated that the magnitude of the outcome effect varied by trial type. Bonferroni–corrected pairwise comparisons showed that the reward outcome effect (win - nothing) was significantly larger than both the loss outcome effect (loss - nothing; Δ = 0.184, SE = 0.058, t = 3.17, p = 0.0047) and the neutral outcome effect (look - nothing; Δ = 0.224, SE = 0.058, t = 3.85, p = 0.0004). The loss and neutral outcome effects did not differ (Δ = 0.040, SE = 0.058, t = 0.69, p = 1.00).

In summary, ‘win/loss/look’ outcomes produced a positive deflection relative to nothing, but this effect was roughly twice as large for win outcomes. These results are summarised in Figure 4B.

Behavioural_Type, classified as NonRandom vs Random on the basis of per-trial-type binomial tests of choice rate against chance (see Methods), did not significantly modulate the within-context outcome contrasts in a three-way Trial_Type × Outcome_Label × Behavioural_Type model (Outcome_Label × Behavioural_Type: F(3, 3836.7) = 1.60, p = 0.19). The reward-linked modulation (win − nothing) was similar in direction and magnitude across behavioural types (NonRandom β = 0.316, p < 0.001; Random β = 0.356, p < 0.001), indicating that the principal outcome effects generalise across behavioural strategies.

Of the four further candidate predictors entered as three-way interactions with Trial_Type and Outcome_Label — Smoking, HAMD, HIT-6, and SHAPS — none survived Bonferroni correction across the five predictors. The Outcome_Label × HAMD term was the only effect to reach unadjusted significance (F(3, 3836.7) = 3.21, p = 0.022; Bonferroni-adjusted p ≈ 0.11).

Higher depression scores were associated with numerically larger outcome contrasts across all three trial types, most pronounced for the neutral look − nothing contrast, which increased from 0.024 at −1 SD to 0.217 at +1 SD HAMD; the corresponding range was 0.27 to 0.40 for the win − nothing contrast and 0.11 to 0.21 for the loss − nothing contrast. Smoking (F(3, 3837.3) = 2.09, p = 0.10), SHAPS (F(3, 3836.7) = 2.34, p = 0.07), and HIT-6 (F(3, 3836.9) = 0.38, p = 0.77) did not reach unadjusted significance. Given the absence of a priori prediction and the bimodal distribution of HAMD scores in our sample, we report the depression-related modulation as exploratory.

Adding clinical covariates (HAMD, HIT-6, SHAPS) as well as Smoking and salience proxies (see Methods for details) did not improve fit over the core model (BIC preferred the core specification), and overall variance explained was modest (marginal R² = 0.022; conditional R² = 0.176).

### Testing for outcome-by-option value interaction following learning

We predicted that efficient learning would produce an interaction between the value of the chosen option (high vs. low) and the outcome (’win’ vs. ’nothing’), since for a high-value option a ’nothing’ outcome would be more surprising, and conversely, a ’win’ would be more surprising for a low-value option. We therefore fitted a linear mixed-effects model (random intercept for subject) to single-trial amplitudes obtained from trials after the first 25 reward trials, averaged over the outcome type effect window (0.317-0.467 s) established in Figure 4A for the eight subjects with gradual or consistent behavioural type.

Given our a priori directional hypothesis that unexpected wins (low-value option chosen) would elicit stronger VTA responses than expected wins (high-value option chosen), we conducted a planned one-sided interaction contrast testing whether (Low:Win − High:Win) − (Low:Nothing − High:Nothing) > 0. This contrast was significant (estimate = 0.316, SE = 0.167, t(676) = 1.89, one-sided p = 0.029), supporting the prediction that the win-related VTA response was modulated by option value in a direction consistent with reward prediction error. Examining the estimated response amplitudes for the four conditions confirmed the expected pattern of effect (Figure 5). Notably, because the interaction contrast is orthogonal to the main effect of outcome type, it was not biased by our selection of the analysis time window based on the latter.

**Figure 5.**
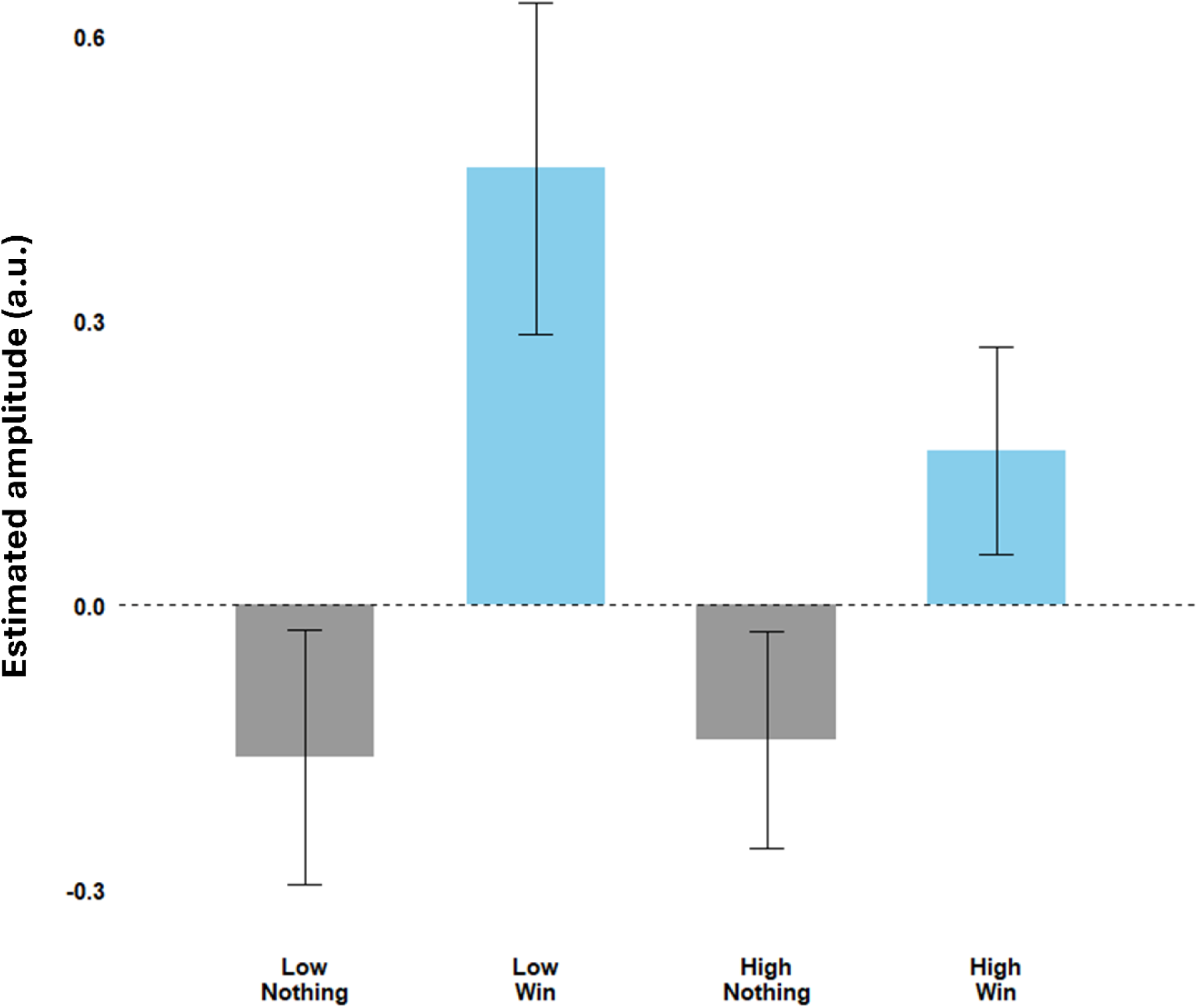
Outcome-by-option value effects following learning. Estimated marginal means (EMMs) of VTA-LFP amplitudes from the outcome window (0.317-0.467 s) after the first 25 reward trials, derived from a linear mixed-effects model (random intercept for subject, N = 8). Bars are grouped by option value (low vs. high probability), with grey = ’nothing’ and blue = ’win’ outcomes. Error bars denote ± 1 SE. A planned one-sided interaction contrast confirmed that the win-evoked VTA response was significantly larger when the chosen option had low expected value (ΔLow − ΔHigh = 0.32, p = 0.029), consistent with reward prediction error encoding.

### Testing for the representation of value

Having established a group-level RPE-consistent signal, we next asked whether trial-by-trial value encoding was detectable at the single-subject level, treating this as a mechanistically informative case analysis rather than a basis for group inference. We, therefore, tested for the value representation in the single subject who displayed Gradual learning consistent with reinforcement learning theory and had clear LFP responses (Subject 3). We performed General Linear Modelling with three different representations of value (value of the chosen, high-value and low-value options), derived from the single learning rate RW-1α model which was the best model for this subject, and confound regressors (see Methods) and used an F-test to test for any value effect in representation-agnostic way.

This revealed two significant clusters around the time of the button press (−0.5 to -0.43 s, p=0.042, peak-level significant at -0.5s p=0.043; -0.1 to 0.033 s, p=0.001 with the peak at - 0.07s, p=0.003). No significant effects were found related to the stimulus presentation and outcome events.

To further interrogate the decision-related effect, we performed post-hoc correlations between the three value regressors and LFP values averaged over the time windows revealed by the F-test.

The results are presented in Supplementary Table S2. They show that for both time windows the correlations are much stronger for the value of the chosen option regressor. This is remarkable given the non-trivial shape of this regressor consisting of sharp transitions between high and low values in adjacent trials (Figure 6A). Figure 6B shows the individual trials of Subject 3 sorted by the value of the chosen option, making it possible to see the origins of these effects. It appears that for low value trials the response is both lower in amplitude and shifted towards the button press time which leads to two clusters of differences first positive (−0.5 to - 0.43s) and then negative (−0.1 to 0.017s).

**Figure 6.**
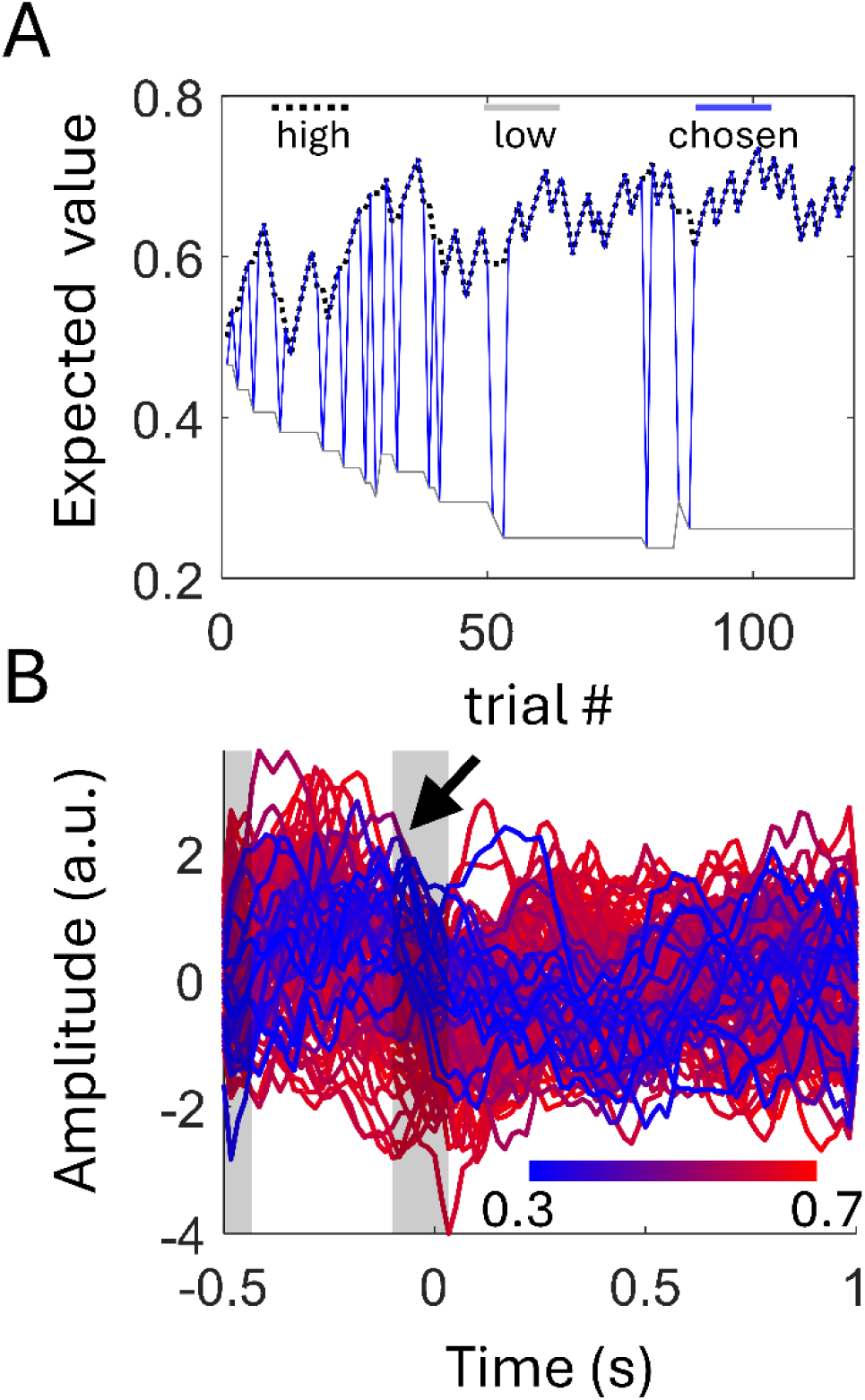
Value correlates in VTA-LFP responses of Subject 3 around the decision event (button press). (**A**) Profile of the ‘value of the chosen option’ regressor across trials (blue line). The black dotted line shows the value of the high value option and the grey solid line – that of the low value option. The blue line alternates between the two depending on the choice. (**B**) Single-trial VTA-LFP responses, colour-coded by the chosen option’s value (blue: lowest; red: highest). Shaded areas indicate significant clusters (cf. Supplementary Table S2).

To rule out the potential contribution of reaction time confound and residual stimulus-locked activity, we tested for the effect of reaction time and found no significant effects in the time window of the effect of value.

We also tested for this effect in consistent choosers (N = 7; Subjects 2, 4, 5, 6, 9, 10 and 12) using value estimates derived from the dual learning rate model (RW–2α), which provided the best fit for these participants. Owing to the small sample size, analyses were restricted to a fixed–effects design combining all the trials with subject–specific regressors. F-test for the effect of value only showed a late cluster (0.817 to 0.85 s, p = 0.022). However, this design also permitted assessment at the individual subject level, revealing a pattern in Subject 12 that closely resembled that observed in Subject 3. Two significant clusters were identified prior to the button press, at −0.50 to −0.45 s (p = 0.004) and −0.117 to −0.033 s (p < 0.001). Supplementary Figure S3 shows the trial–wise analysis for Subject 12, which closely resembles that shown in Figure 6, albeit with fewer low–value trials. This difference reflects the fact that Subject 12 only explored the low–value option occasionally during the second half of the experiment.

## Discussion

Our study provides the first direct electrophysiological evidence that the human VTA encodes reward-related signals during motivated decision-making, bridging decades of animal electrophysiology with human neuroscience. Recording directly from this otherwise inaccessible dopamine-rich structure, made possible by a rare neurosurgical opportunity, we found that VTA population-level responses were significantly larger for rewarding outcomes than for either loss or neutral outcomes. In contrast, there was no significant difference between responses to losses and neutral outcomes, indicating that the human VTA is selectively sensitive to reward. In behavioural terms, subjects also tended to respond faster on reward trials than on loss trials, consistent with heightened motivational engagement when potential rewards were at stake. Together, these findings suggest that the human midbrain, like in other species, is preferentially tuned to detect and respond to rewarding events.

The observation that participants who developed a preference for the high–value option also showed enhanced responses to low–probability wins suggests that VTA-LFP activity reflects more than a simple encoding of outcome. Instead, these responses appear to integrate outcome and expected value, consistent with reward prediction error signalling as formalised in reinforcement learning theory.

Our results confirm in humans a neural dynamic long established in animal models of dopamine function. Decades of work in rodents and primates have shown that VTA dopaminergic neurons fire strongly to unpredicted rewards but show little or no excitation for aversive or omitted outcomes (signalling a positive RPE). This asymmetric coding – primarily representing reward rather than punishment – was evident in our human VTA recordings, aligning with classic theories of reinforcement learning ^2,3,23^. Moreover, our findings extend prior indirect and adjacent evidence from humans. Functional MRI studies have inferred that BOLD activity in the midbrain scales with RPE magnitude (e.g. ^24,25^) and EEG studies link frontal signals like the feedback-related negativity to dopaminergic error processing ^26^. Perhaps most relevant, intraoperative single-unit recordings from the neighbouring substantia nigra in patients showed neurons that responded more strongly to unexpected rewards than to losses ^14^. We now bridge these observations by demonstrating directly that population-level activity in the human VTA carries a similar reward-predictive signal. Notably, animal studies have reported that also LFP signals in the VTA encode reward information and adapt to changing cue–reward contingencies ^27–29^. Our human LFP results echo this cross-species consistency, providing a crucial translational link between animal electrophysiology and human reward processing.

These recordings were made possible by a rare clinical opportunity - leveraging DBS electrodes implanted in the VTA of patients with severe chronic cluster headache. This unique context allowed us to probe human midbrain physiology that is otherwise inaccessible. However, task engagement varied, limiting interpretation. Three subjects experienced acute cluster headache attacks during the experiment, which likely impaired their focus and learning and all the subjects were tested just a few days after invasive brain surgery done under general anaesthesia and after sleeping on a noisy hospital ward. As a result, only nine of fourteen subjects developed a clear preference for the high-reward option. Of these, just two showed gradual, trial-by-trial learning predicted by classical reinforcement learning models. Most learners adopted an immediate ’win-stay’ strategy, which is itself formally captured by the dual learning rate RW-2α model: very high reward learning rates combined with near-zero non-reward learning rates produce exactly this pattern of rapid commitment to the high-value option. Far from invalidating the RL framework, this asymmetry demonstrates that participants were efficiently exploiting reward contingencies, albeit with limited exploration of the low-probability alternative.

Crucially, elevated VTA response to wins compared to losses and neutral outcomes was not dependent on behavioural strategy and was also observed in subjects with random responses. This indicates that VTA is specifically tuned to rewards, independently of the behavioural context.

In a single gradual learner with clear VTA responses (Subject 3), enabling a full test of reinforcement learning predictions, VTA evoked activity tracked the trial–by–trial value of the chosen option. Although a single case cannot support group–level inference, the finding is notable for several reasons. First, the chosen value was a rapidly fluctuating, non–trivial regressor during early learning, and its capture suggests sensitivity of the VTA signal to ongoing learning. Second, the effect persisted after controlling for fatigue, time–on–task and reaction time, arguing against confounds such as arousal, recording drift or stimulus–locked responses. Third, the timing was theoretically plausible: value modulation emerged before the button press during decision formation, and on low–value choices the response was delayed and attenuated, consistent with weaker reward expectation. Fourth, the effect was time–locked to the button press rather than the stimulus, supporting a decision–related interpretation. Fifth, a similar pattern was observed in another subject who occasionally explored the low–value option despite predominantly choosing the high–value option. Together, these observations suggest that during active, gradual learning, human VTA activity can encode expected value in real time. We interpret this cautiously given the limited sample, but it aligns with core predictions of temporal–difference learning in dopamine circuits^30^.

In summary, this study provides the first direct evidence that human midbrain activity exhibits key signatures of reward processing and reinforcement learning previously demonstrated in animals. By recording electrophysiological signals during decision-making, we show that this dopamine-rich region responds strongly to rewarding outcomes and, under effective learning, could reflect expected value before choice. These findings validate the translational relevance of animal models and offer a foundation for exploring how reward signals vary across behavioural contexts and clinical populations. Clinically, these findings carry direct relevance beyond cluster headache. Disruption of VTA reward signalling is implicated in depression, addiction, and Parkinson’s disease; conditions increasingly targeted by neuromodulation therapies. Establishing that human VTA population activity carries identifiable reward and value signals provides a principled electrophysiological target for understanding how DBS and related interventions alter motivational processing and may ultimately inform biomarker development for treatment response. More broadly, this work demonstrates that rare neurosurgical access, combined with computational modelling, can test longstanding theoretical predictions about human reward circuitry that are otherwise inaccessible to direct investigation.

## Materials and methods

### Subjects and surgery

The study received approval from the East Midlands – Derby National Health Service Research Ethics Committee, and all subjects provided written informed consent. Fourteen subjects (nine male, age 46±10) undergoing DBS surgery for chronic cluster headache treatment were recruited. No prior human studies provided effect size estimates for power calculations. We therefore used a sample size comparable to previous intracranial DBS studies (10–20 participants), constrained by the rarity of eligible patients and the need to complete recruitment over several years. Hamilton Depression Rating Scale 21-item version (HAM-D 21) and Headache Impact Test-6 (HIT-6) scores were collected on the day of the experiment, except for subject 14, who completed the questionnaires several months later. The HAM-D 21 is a widely used clinical assessment tool measuring the severity of depressive symptoms in individuals with mood disorders. The HIT-6 assesses the impact of headaches on daily life and functioning. The study flow diagram is shown in Figure 1A, and subject details and scores are summarised in Supplementary Table S1.

VTA electrodes were implanted via MRI-guided and image-verified stereotactic surgery under general anaesthesia. Eight subjects had bilateral leads: the rest unilateral. Figure 1B shows the lead locations reconstructed with Lead-DBS (https://www.lead-dbs.org/, ^31^). After surgery, the VTA leads were left externalised for several days. The externalisation allowed for recording of VTA-LFPs, both during periods of rest and while the subject performed an instrumental learning task.

### Task

The instrumental learning task used was a modified version of a static probabilistic reinforcement learning task that has been used extensively in previous studies ^32,33^, involving three trial types with rewarding, neutral, and aversive outcomes linked to distinct fractal image pairs. Each trial type had its specific image pair, with associations randomized across subjects. ’Win’ trials could result in ’win’ or ’nothing’ while ’loss’ trials could lead to ’loss’ or nothing’. A neutral condition had ’look’ or ’nothing’ outcomes. Unbeknown to the subjects, one fractal in each win/loss pair had a high probability (0.7) of the corresponding outcome and the other a low probability (0.3). Subjects aimed to maximize ’vouchers’ won and minimize losses through trial and error, although no real payments were involved. Each trial followed a structured sequence. The trial began with the presentation of a fractal pair for 1.5 seconds, during which subjects had to make a response. If no response was made within this 1.5-second window, the trial was aborted and excluded from further analysis. Following a response, there was a variable delay before the outcome appeared, ranging between 1.6 and 3 seconds. The outcome label (‘You win’, ‘You Lost’, ‘Look’ or ‘Nothing’) was displayed for 2 seconds. Following the outcome label presentation, an inter-trial interval (ITI) of 2 seconds occurred before the next trial. For Subject 1, the interval between button presses and outcome label presentation ranged from 4 to 6 seconds, while for Subject 2, it ranged from 3.5 to 6 seconds. In subsequent subjects, this interval was standardized to 1.6 to 3 seconds. Additionally, the ITI was initially jittered in the first two subjects but was later fixed at 2 seconds to save time. A summary of the task is shown in Figure 2A.

### Behavioural analysis

#### Model-free behavioural validation

To provide a model-free assessment of learning, we fitted mixed-effects logistic regression models (GLMM^34^) with a random intercept for subject, testing whether the group-level probability of choosing the better option exceeded chance (50%) for reward and loss trials (one-sided tests), and whether any systematic preference was present in neutral trials (two-sided test). Individual-level binomial tests were also conducted to identify subjects performing significantly above chance.

#### Model specification and comparison

To characterise individual choice behaviour in the reward trials, we compared three computational models of increasing complexity using the Bayesian Information Criterion (BIC). The first was a Null model assuming random responding (P(choice) = 0.5 on every trial), serving as a baseline. The second was a single-learning-rate Rescorla–Wagner model ^35^ (RW-1α) with two free parameters: a common learning rate (α) applied to all outcomes and an inverse temperature (β) governing choice determinism. The third was the dual-learning-rate Rescorla–Wagner model (RW-2α) with three free parameters: separate learning rates for rewarding (α_rew_) and non-rewarding (α_nothing_), and an inverse temperature (β). This model could better accommodate cases with a strong preference for one of the options. A similar dual-learning-rate RL model has been applied in human populations by ^36^ to explain differential sensitivity to reward and punishment in Parkinson’s disease. The individual parameters were drawn from group-level distributions defined by hyperparameters for the mean (μ) and standard deviation (σ), ensuring stabilized estimates through partial pooling.

On each trial, expected values (EV) for the chosen and unchosen fractal images are updated as follows:

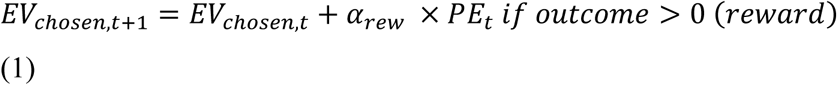

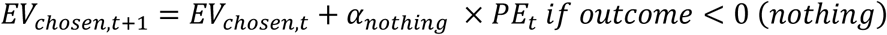

where the prediction error (𝑃𝐸_𝑡_) is calculated as

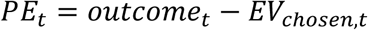

The unchosen fractal’s value remains unchanged in this model, as no fictive updates are applied. The probability of choosing a given fractal is determined using a softmax function:

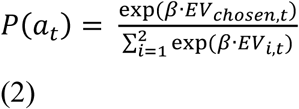

Here, i indexes the two options (e.g., fractal images) available on a given trial, ensuring that the denominator sums the exponentiated expected values of all options, normalising the probabilities. For the RW-1α model α_rew_ and α_nothing_ were set to be equal.

For each subject, BIC was computed as BIC = −2 × log-likelihood + k × ln(T), where k is the number of free parameters and T is the number of reward trials. The model with the lowest BIC was selected as best-fitting for each subject. Based on the winning model and examination of individual choice trajectories, each subject was assigned a behavioural type: Random (R) — the Null model yielded the lowest BIC, indicating no evidence of contingency-sensitive responding; Gradual (G) — the RW-1α model won and the subject initially explored both options before converging on the high-value choice; Consistent (C) — the RW-2α model won and the subject predominantly chose one option with minimal exploration of the alternative.

#### Model fitting and estimation

We implemented the model in a hierarchical Bayesian framework using Markov Chain Monte Carlo (MCMC) sampling in Stan (https://mc-stan.org/). The model was fitted with 6,000 iterations per chain, including a 3,000-iteration warmup period, across 4 chains, with a random seed set for reproducibility. The group-level parameters for learning rates and decision sensitivity were modelled as normally distributed. Learning rates were constrained to the range [0,1], while decision sensitivity (β) was scaled to the range [0,10] using an inverse probit transformation followed by a multiplication factor. Model fit was evaluated based on the log-likelihood of the observed choices, and posterior predictive checks were performed to ensure that the model accurately reproduced observed choice behaviour. Convergence diagnostics, including the Gelman-Rubin statistic (𝑅^), confirmed that all parameters achieved adequate convergence.

#### Electrophysiological data acquisition

The LFP recording in Subject 1 was done using TMSi Porti 32-channel EEG system (TMSi, Oldenzaal, The Netherlands), referenced to the average across channels, sampled at 2048Hz. For the remaining 13 subjects LFP recordings were performed using BrainAmp MR+ EEG system (Brain Products GmbH, Gilching Germany), referenced to the right mastoid, sampled at 2500Hz.

#### Electrophysiological data analysis

The data were analysed using Statistical Parametric Mapping toolbox (SPM, http://www.fil.ion.ucl.ac.uk/spm/, ^37^). Following conversion to SPM format, the LFP recordings were converted to a bipolar montage between adjacent contacts (3 channels per hemisphere). The data were downsampled to 300Hz. Digital filtering was applied to remove frequency components below 0.5Hz (Butterworth zero phase 5^th^ order filter). The data were then epoched into segments from -0.5s to 1s relative to the events of interest. Trials containing outliers exceeding 5 standard deviations, computed for each channel separately across the whole recording, were excluded from analysis.

For time-frequency analysis, Morlet wavelet transform ^38^ was used with a fixed time window of 50 ms for frequencies between 1 and 100Hz with 2.5Hz spacing. The time axis was subsampled by a factor of 4 and cropped between -0.45 to 0.95s to remove edge effects. The power was conditioned by taking the log prior to statistical analysis. Phase information was saved separately and used for computing inter-trial coherence (ITC) ^39,40^

#### Analysis of evoked and induced responses

Trial-by-trial LFP waveforms and time-frequency images for each subject were entered into a regression model (SPM design matrix). Depending on the analysis, the design matrix could include only the mean term or the mean term and the effect of outcome label (win/loss/look vs. nothing). For reward trials of the subjects for whom the choice value could be computed (see Results), we used a more complicated design matrix with the following regressors:

Confound regressors

1. Constant regressor modelling the intercept
2. Binary regressor modelling the outcome label (‘win’ or ‘nothing’)
3. Time since the beginning of the experiment
4. Trial number in the current recording block
5. Reaction time

Regressor of interest

6. Value of the chosen option

The design matrix was inverted using Matlab’s *pinv* function and the resulting weights were applied to the trial data in time and time-frequency domains to generate waveforms or images of regression coefficients respectively (see spm_eeg_contrast function in SPM). ITC images were computed for the mean response to each event across all trial types with the aim of distinguishing genuine induced responses from time-frequency representations of the evoked response.

The waveforms were low-pass filtered at 20 Hz, down-sampled to 60 Hz, averaged across all the LFP channels, baseline-corrected, and normalised by a subject-specific scaling factor to account for differential gain across subjects. The scaling factor was the epoched LFP’s standard deviation, computed across all channels, events and conditions after rejecting bad trials.

Time-frequency images of GLM coefficients and ITC were averaged across channels and baseline corrected (−0.45 to -0.05s).

#### Linear mixed-effects analysis

To examine how behavioural and clinical variables modulate electrophysiological responses, we fitted linear mixed-effects (LMM) models in R (https://www.r-project.org/) using the lmerTest/lme4 framework ^41^. Trial-wise VTA-LFP amplitude, averaged over the time window where a significant effect of outcome label was detected, was the dependent variable. Outcomes were analysed as the observed Outcome_Label factor (levels: win, loss, look, nothing) within Trial_Type (Reward, Loss, Neutral). The core fixed-effects structure was Trial_Type × Outcome_Label, with by-subject random intercepts. This core specification was retained as the reference model for all subsequent analyses.

We then tested five candidate predictors of the within-context outcome contrasts, each entered as a three-way interaction with Trial_Type and Outcome_Label in a separate model:

Behavioural_Type (NonRandom vs Random; derived from per-trial-type binomial tests of choice rate against chance, see Table 1),

Smoking (yes/no, to test for the possible effects of nicotine on VTA responses⁴¹),

HAMD (Hamilton Depression Rating Scale),

HIT-6 (Headache Impact Test-6),

SHAPS (Snaith-Hamilton Pleasure Scale).

For each predictor, the principal test was the Outcome_Label × predictor term in a Type-III ANOVA. The Behavioural_Type reference level was Random, so the NonRandom coefficient is interpretable as the effect of being a learner. All continuous covariates (HAMD, HIT-6, SHAPS) were z-scored within sample. Missing SHAPS values were imputed by group mean within Smoking strata, as pre-specified.

As an auxiliary check on the parsimony of the core specification, we compared the bare core to additive extensions including Smoking; the full clinical/demographic block (Smoking + Age + HAMD + HIT-6 + SHAPS); and the clinical block plus trial-wise salience covariates: log reaction time (logRT), expected value of the chosen option (EV_chosen), elapsed block time (BlockTime), and the previous-trial outcome (PrevOutcome). Model selection was based on the Bayesian Information Criterion (BIC), where lower BIC reflects a better fit–complexity trade-off. Random effects comprised by-subject intercepts; Trial_Type random slopes were included where estimable. Models showing convergence warnings or singular fits were simplified to random intercept only. Post-hoc estimates used estimated marginal means (EMMs). Planned within-context contrasts (Reward: win−nothing; Loss: loss−nothing; Neutral: look−nothing) were evaluated with Šidák adjustment for multiplicity and Satterthwaite degrees of freedom; pairwise comparisons across contexts used Bonferroni correction. All predictor tests were verified against Kenward–Roger degrees of freedom as a sensitivity check. We report β (SE), df, t, p, and marginal/conditional R² (variance explained by fixed effects alone vs. by fixed + random effects).

#### Testing for outcome-by-value effects following learning

To examine whether outcome responses in reward trials were influenced by learned expectations, we analysed single-trial LFP amplitudes after the first 25 reward trials, corresponding to the post-learning phase. For each subject, mean LFP amplitudes were extracted within the significant outcome-related cluster (0.283–0.500 s; see Analysis of the response to outcome). Each trial was then categorised according to the value of the chosen option (low or high probability) and outcome label (‘win’ or ‘nothing’). Only subjects who demonstrated gradual or consistent learning behaviour were included (N = 8).

A linear mixed-effects model was fitted in R (lme4 and lmerTest packages) with LFP amplitude as the dependent variable, Value, Outcome, and their interaction as fixed effects, and Subject as a random intercept term. Type III ANOVA with Satterthwaite approximations was used to test main and interaction effects. Planned pairwise comparisons were conducted using estimated marginal means (emmeans package) with Bonferroni correction across the four condition means.

## Data availability

The data and code that support the findings of this study will be made openly available via *figshare.com* at http://doi.org/10.6084/m9.figshare.31029091.

## Acknowledgements

In preparation of the manuscript large language models implemented in ChatGPT and MS Copilot were used for language editing of human-written text. Their outputs were checked for accuracy by the authors.

## Author contributions

Conceptualisation – DS, VL

Electrophysiological data collection – DS, VL

Clinical support, clinical imaging and clinical data collection – MSM, SL, LS, DS, LZ, HA

Data analysis – AR, VL, DS, JPR

Funding Acquisition, Supervision – UV, DS, VL

Writing – original draft – AR, DS, JPR, VL

Writing – review & editing – UV, HA, LZ, MSM, SL, LS

## Supplementary material

Supplementary material is available online.

