## Supplementary material for "Electrophysiological Correlates of Reward Processing in the Human Ventral Tegmental Area"

**Supplementary Table S1. Subject Information.**

| Subject ID | Gender | Age | HIT-6 | HAM-D | Side Implanted |
|---|---|---|---|---|---|
| 1 | M | 54 | 72 | 15 | Left |
| 2* | M | 46 | 76 | 28 | Both |
| 3* | F | 55 | 78 | 5 | Right |
| 4* | F | 40 | 66 | 9 | Both |
| 5* | M | 58 | 71 | 7 | Both |
| 6* | M | 38 | 65 | 8 | Left |
| 7 | M | 60 | 71 | 6 | Right |
| 8 | F | 37 | 71 | 26 | Both |
| 9* | M | 45 | 66 | 0 | Left |
| 10* | M | 62 | 66 | 4 | Right |
| 11 | M | 30 | 72 | 30 | Both |
| 12* | M | 49 | 67 | 15 | Left |
| 13 | M | 36 | 70 | 20 | Both |
| 14 | M | 41 | 64 | 0 | Right |

*The subjects indicated by an asterisk were included in the analysis of the expected value and reward prediction error effects (See Results). Hamilton Depression Rating Scale 21-item version (HAM-D 21) and Headache Impact Test-6 (HIT-6) scores were collected on the day of the experiment, except for subject 14, who completed the questionnaires several months later. The HAM-D 21 is a widely used clinical assessment tool measuring the severity of depressive symptoms in individuals with mood disorders. The HIT-6 assesses the impact of headaches on daily life and functioning.

### Individual choice behaviour for Subjects

**Supplementary Figure S1. Individual choice behaviour for Subjects 1-14 across all three trial types.** Each panel shows trial-by-trial choice data for reward (top), loss (middle), and neutral (bottom) conditions. Faded dots indicate raw choices on each trial (1 = chose better option, 0 = did not). The dashed line shows the 10-trial rolling average, reflecting local choice trends. The solid line shows the cumulative proportion of better-option choices across all preceding trials. The horizontal dashed grey line indicates chance performance (50%).

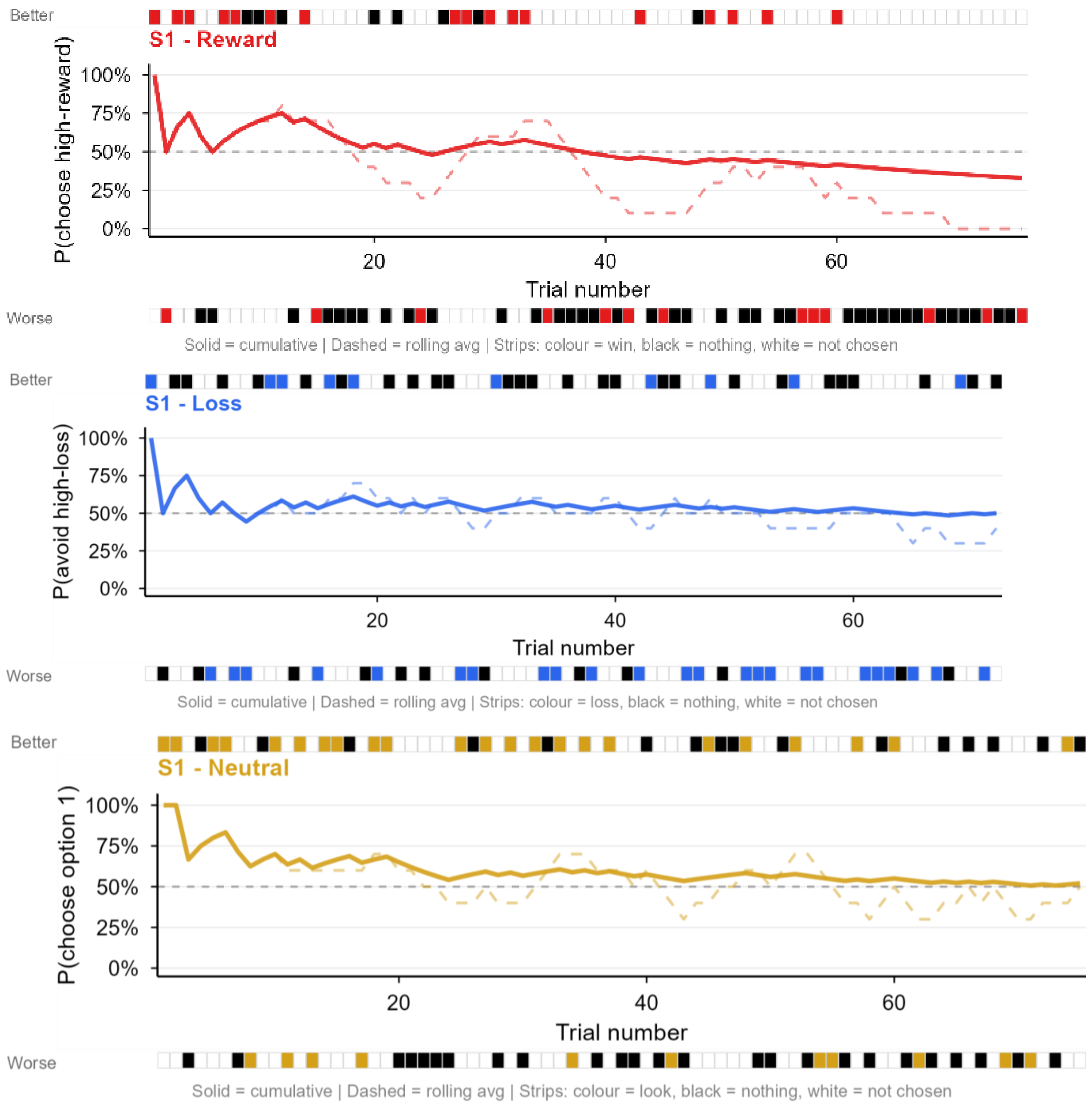

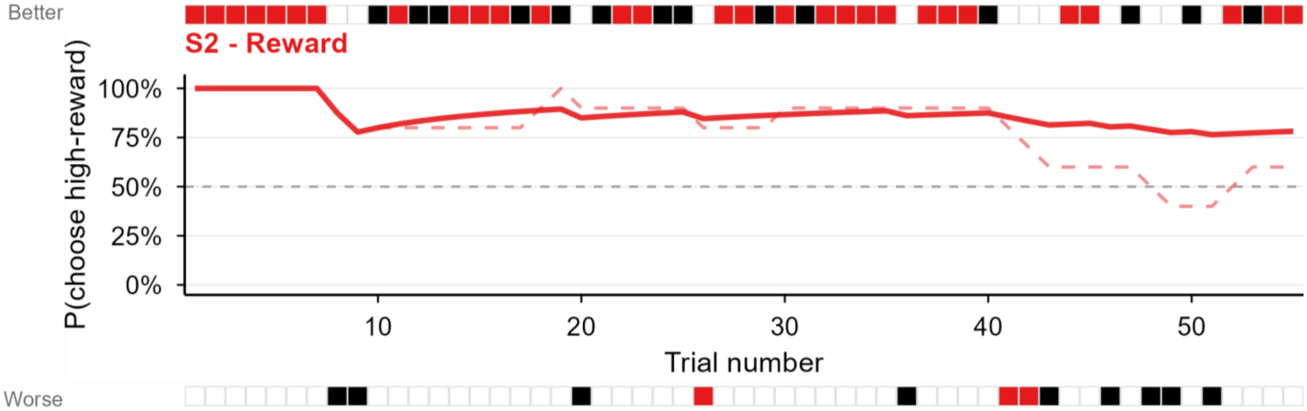


Solid = cumulative | Dashed = rolling avg | Strips: colour = win, black = nothing, white = not chosen

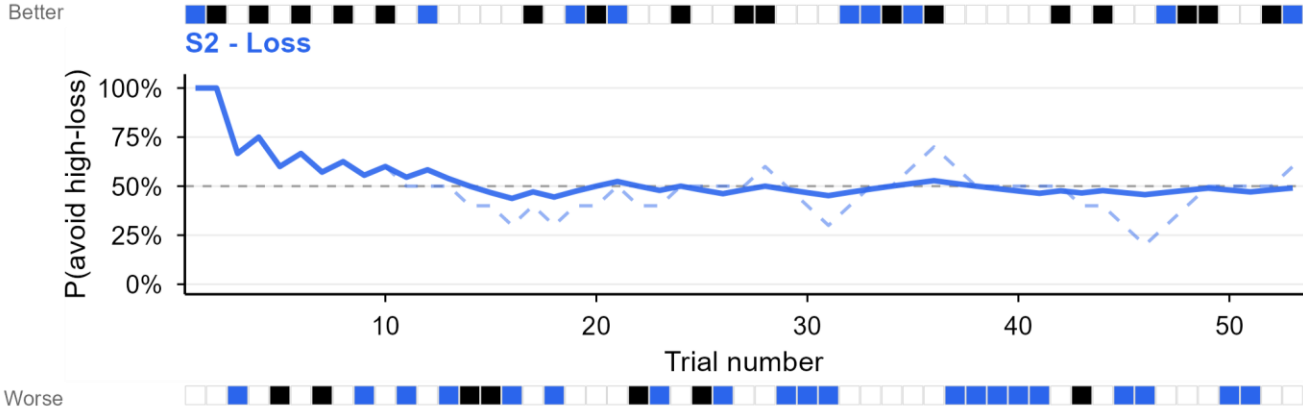


Solid = cumulative | Dashed = rolling avg | Strips: colour = loss, black = nothing, white = not chosen

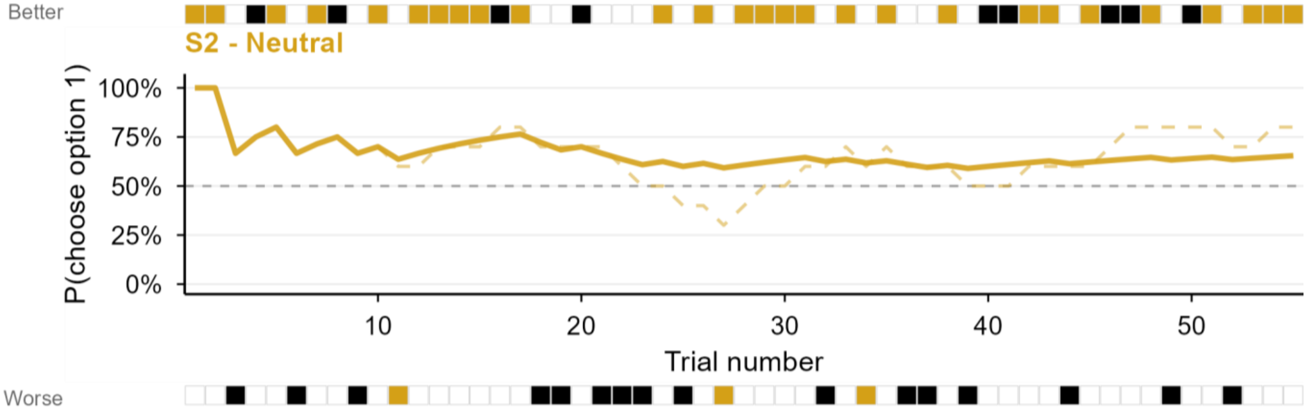


Solid = cumulative | Dashed = rolling avg | Strips: colour = look, black = nothing, white = not chosen

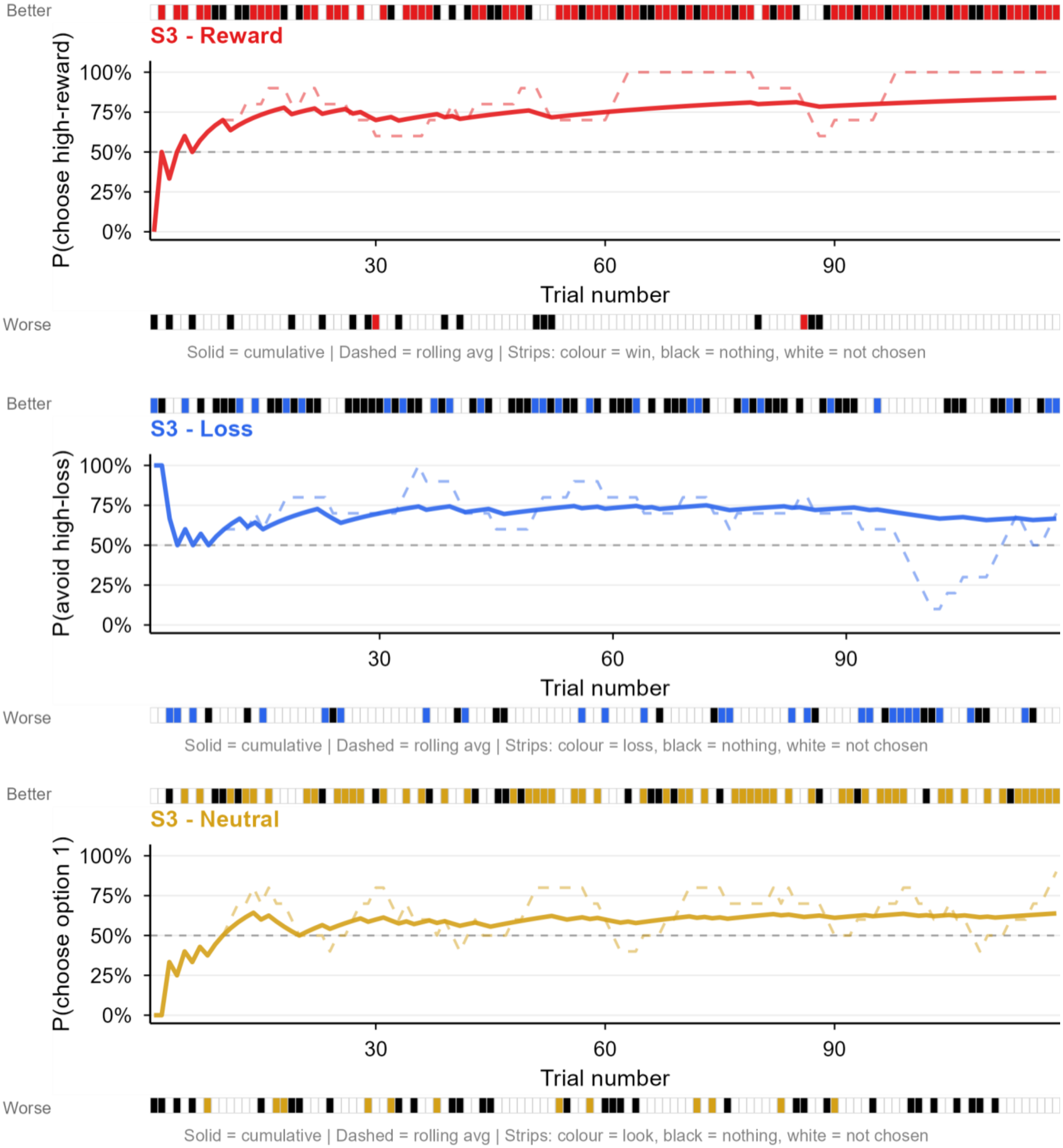
Better
S3 - Reward
P(choose high-reward)
100%
75%
50%
25%
0%
30
60
90
Trial number
Worse
Solid = cumulative | Dashed = rolling avg | Strips: colour = win, black = nothing, white = not chosen
Better
S3 - Loss
P(avoid high-loss)
100%
75%
50%
25%
0%
30
60
90
Trial number
Worse
Solid = cumulative | Dashed = rolling avg | Strips: colour = loss, black = nothing, white = not chosen
Better
S3 - Neutral
P(choose option 1)
100%
75%
50%
25%
0%
30
60
90
Trial number
Worse
Solid = cumulative | Dashed = rolling avg | Strips: colour = look, black = nothing, white = not chosen

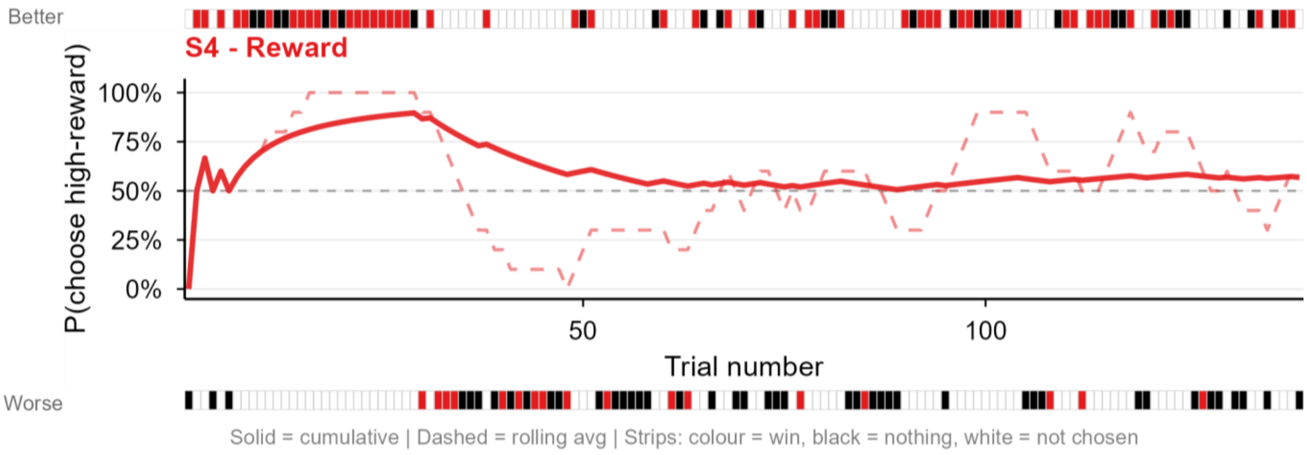

Better
S4 - Reward
P(choose high-reward)
100%
75%
50%
25%
0%
50
100
Trial number
Worse
Solid = cumulative | Dashed = rolling avg | Strips: colour = win, black = nothing, white = not chosen


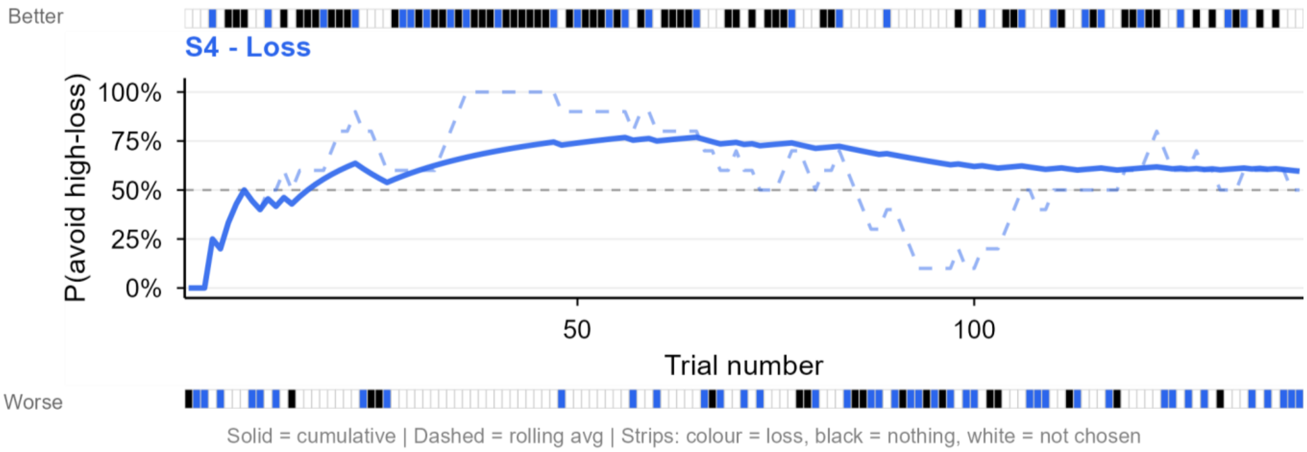

Better
S4 - Loss
P(avoid high-loss)
100%
75%
50%
25%
0%
50
100
Trial number
Worse
Solid = cumulative | Dashed = rolling avg | Strips: colour = loss, black = nothing, white = not chosen


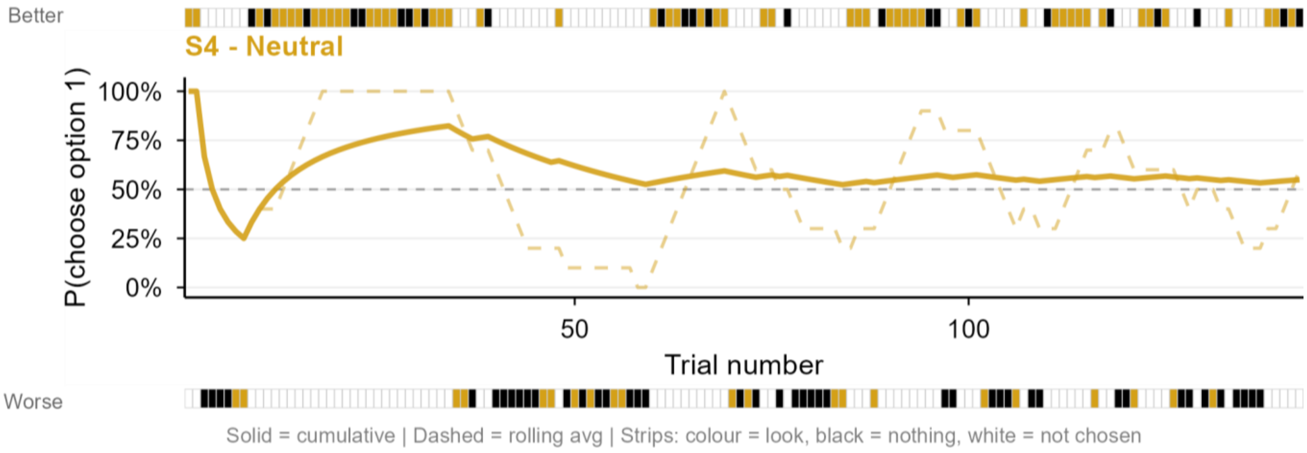

Better
S4 - Neutral
P(choose option 1)
100%
75%
50%
25%
0%
50
100
Trial number
Worse
Solid = cumulative | Dashed = rolling avg | Strips: colour = look, black = nothing, white = not chosen

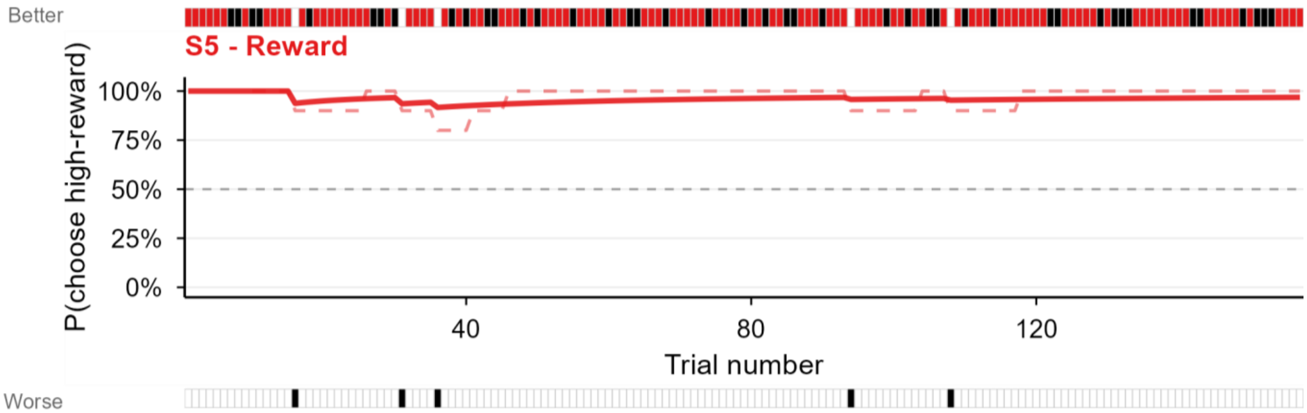


Solid = cumulative | Dashed = rolling avg | Strips: colour = win, black = nothing, white = not chosen

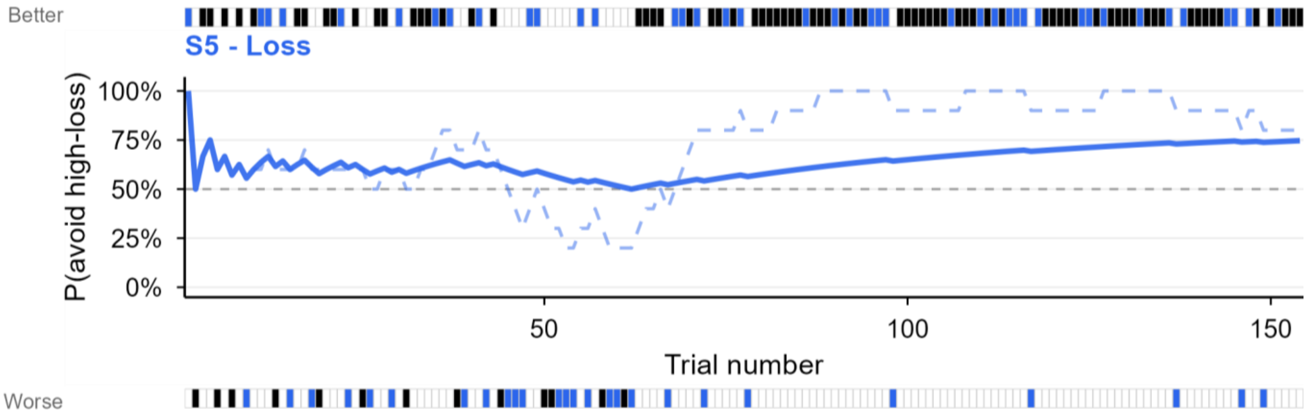


Solid = cumulative | Dashed = rolling avg | Strips: colour = loss, black = nothing, white = not chosen

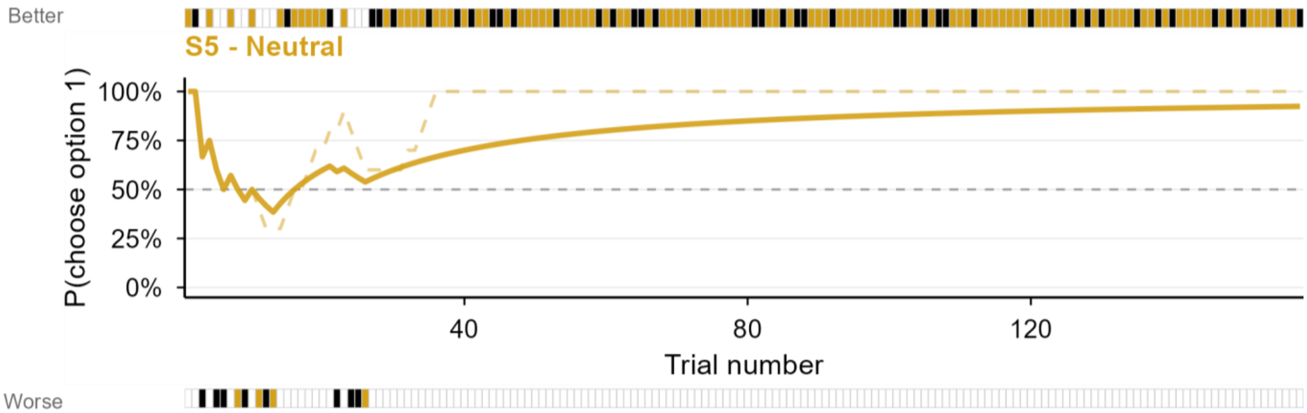


Solid = cumulative | Dashed = rolling avg | Strips: colour = look, black = nothing, white = not chosen

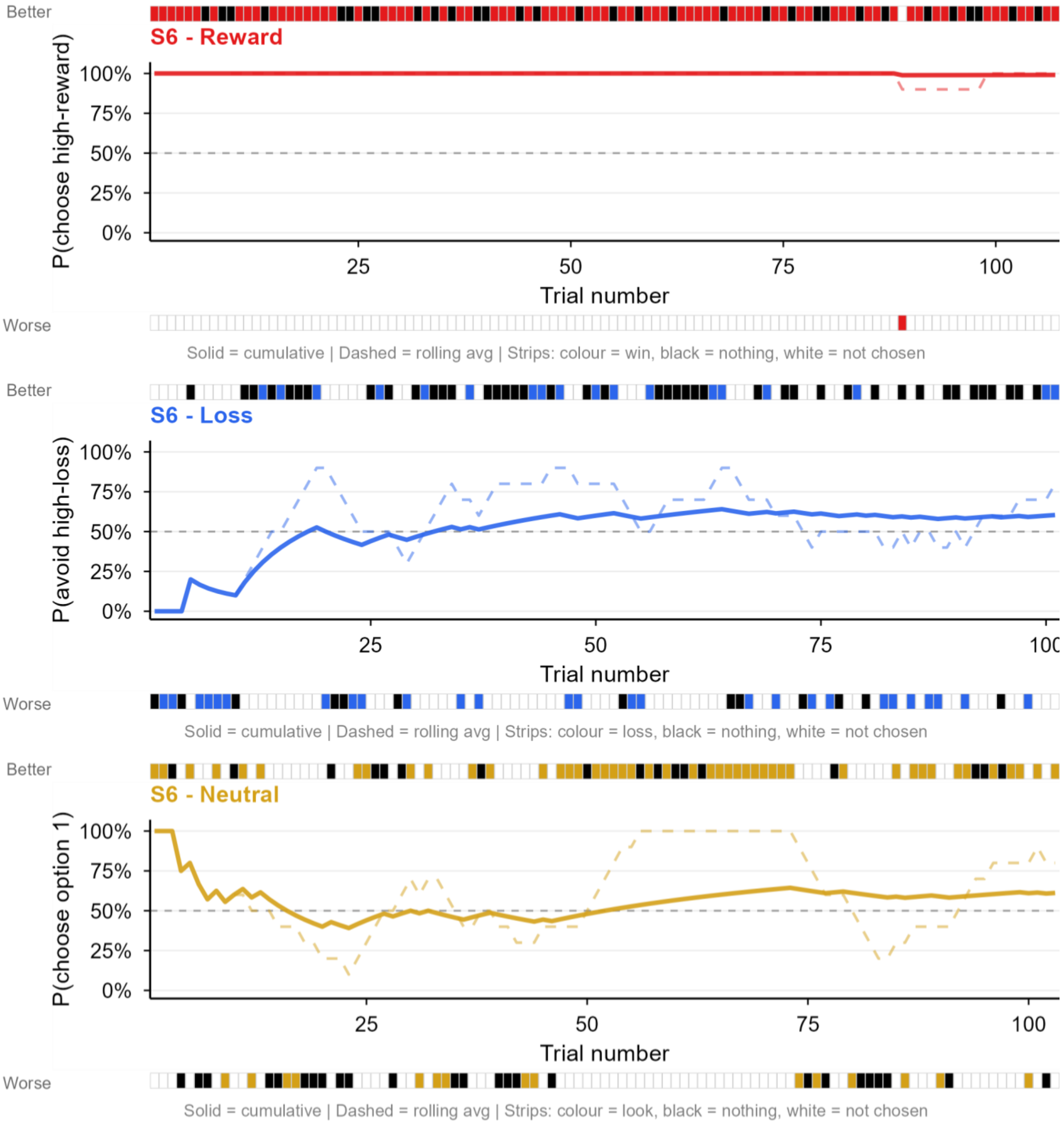

Better
S6 - Reward
P(choose high-reward)
100%
75%
50%
25%
0%
25
50
75
100
Trial number
Worse
Solid = cumulative | Dashed = rolling avg | Strips: colour = win, black = nothing, white = not chosen
Better
S6 - Loss
P(avoid high-loss)
100%
75%
50%
25%
0%
25
50
75
100
Trial number
Worse
Solid = cumulative | Dashed = rolling avg | Strips: colour = loss, black = nothing, white = not chosen
Better
S6 - Neutral
P(choose option 1)
100%
75%
50%
25%
0%
25
50
75
100
Trial number
Worse
Solid = cumulative | Dashed = rolling avg | Strips: colour = look, black = nothing, white = not chosen

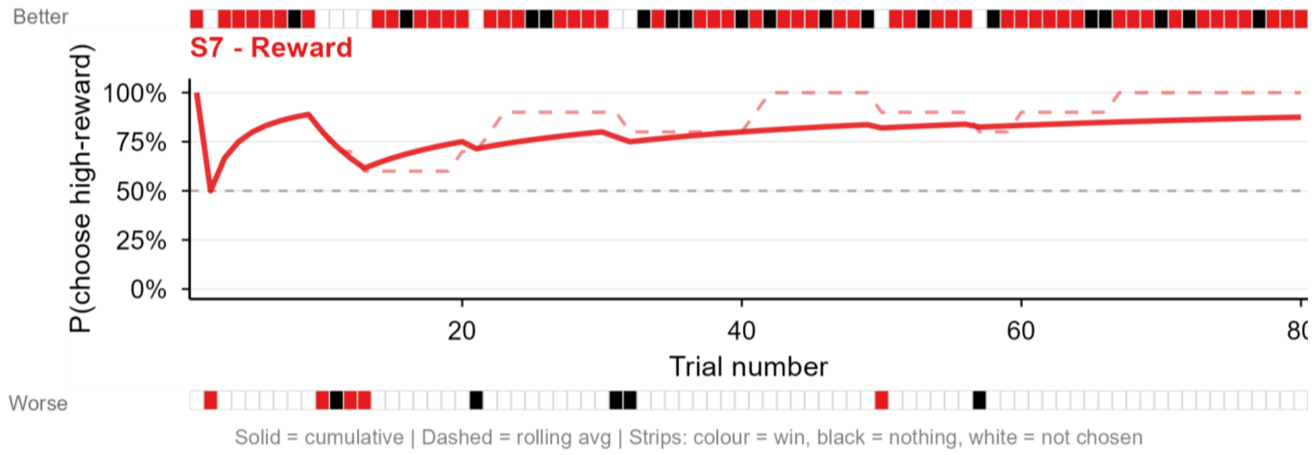
Better
S7 - Reward
P(choose high-reward)
100%
75%
50%
25%
0%
20
40
60
80
Trial number
Worse
Solid = cumulative | Dashed = rolling avg | Strips: colour = win, black = nothing, white = not chosen

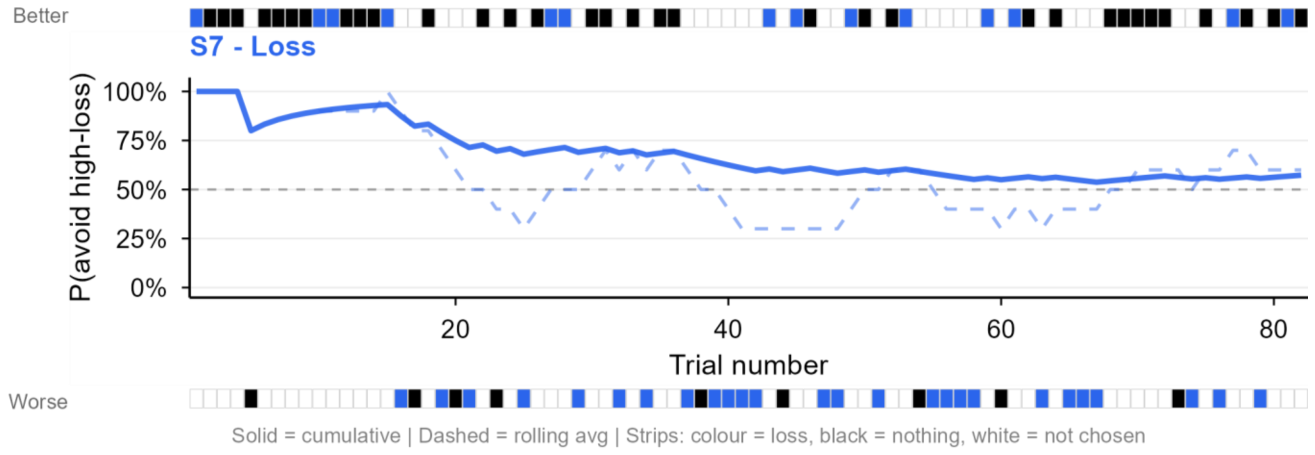
Better
S7 - Loss
P(avoid high-loss)
100%
75%
50%
25%
0%
20
40
60
80
Trial number
Worse
Solid = cumulative | Dashed = rolling avg | Strips: colour = loss, black = nothing, white = not chosen

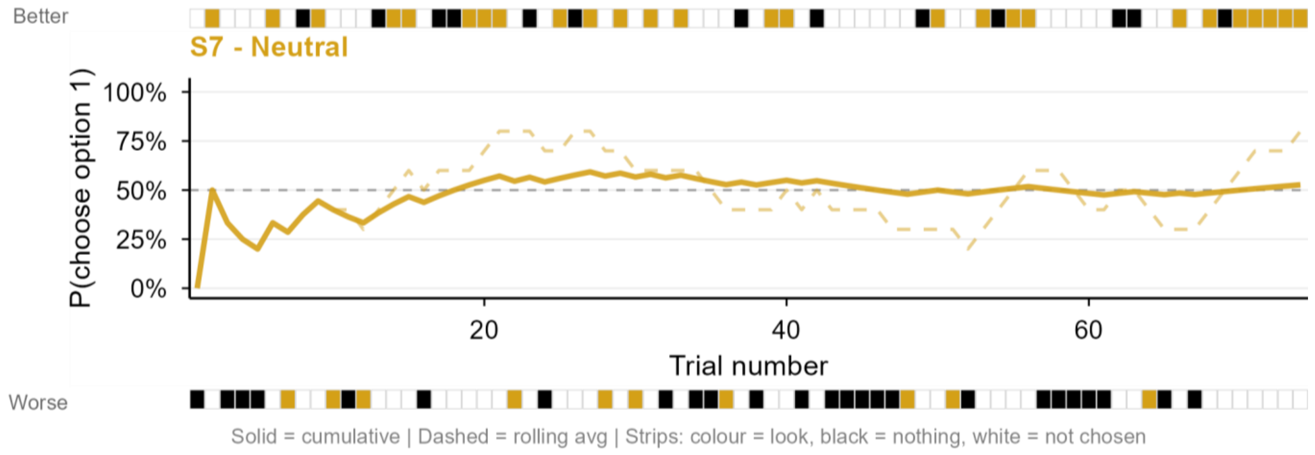
Better
S7 - Neutral
P(choose option 1)
100%
75%
50%
25%
0%
20
40
60
Trial number
Worse
Solid = cumulative | Dashed = rolling avg | Strips: colour = look, black = nothing, white = not chosen

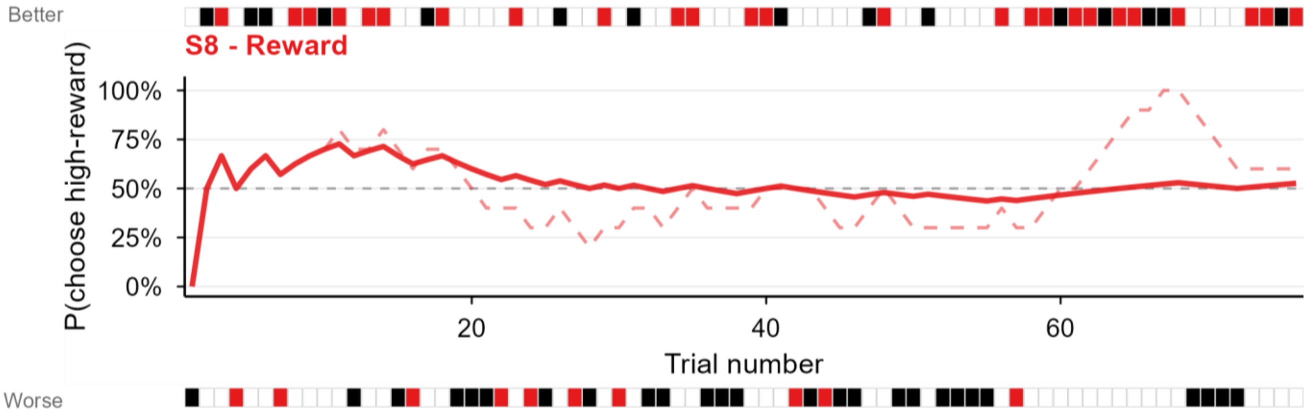


Solid = cumulative | Dashed = rolling avg | Strips: colour = win, black = nothing, white = not chosen

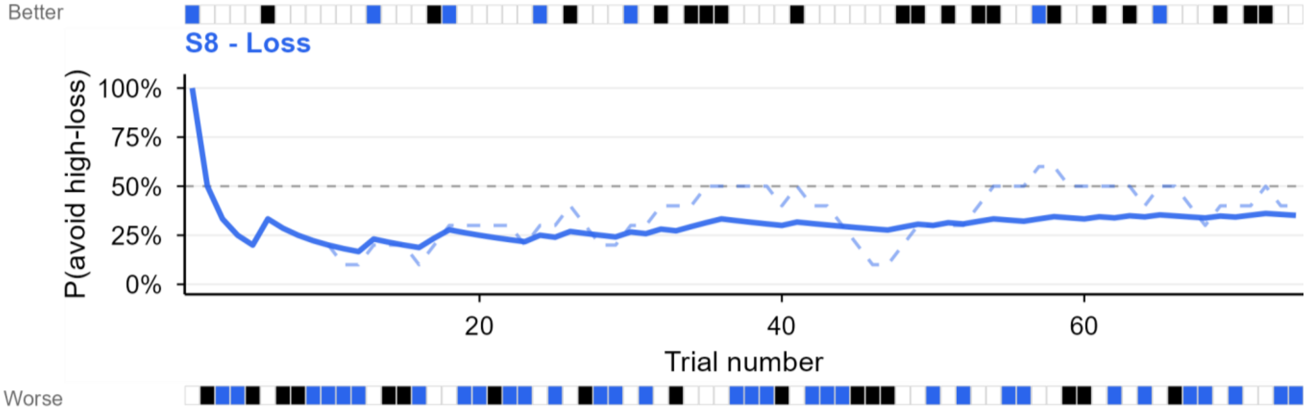


Solid = cumulative | Dashed = rolling avg | Strips: colour = loss, black = nothing, white = not chosen

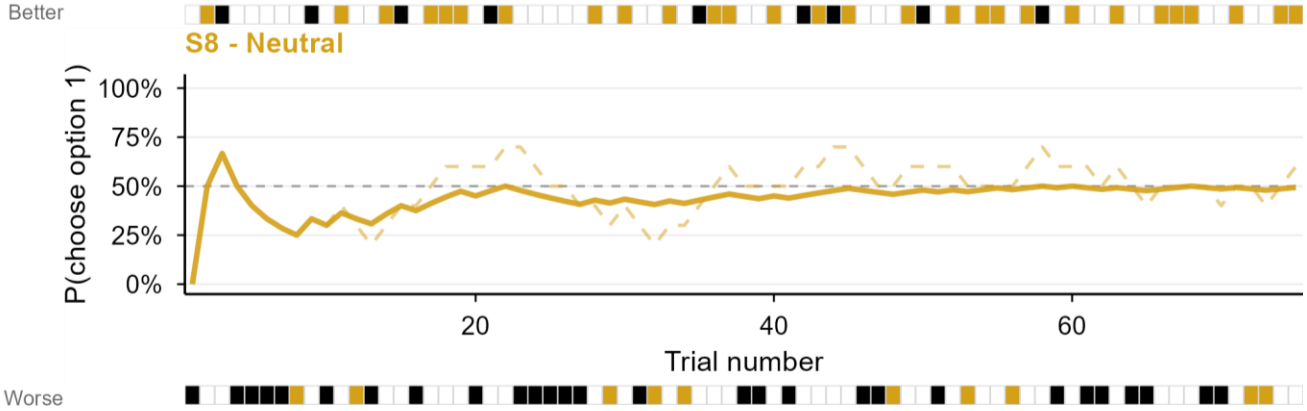


Solid = cumulative | Dashed = rolling avg | Strips: colour = look, black = nothing, white = not chosen

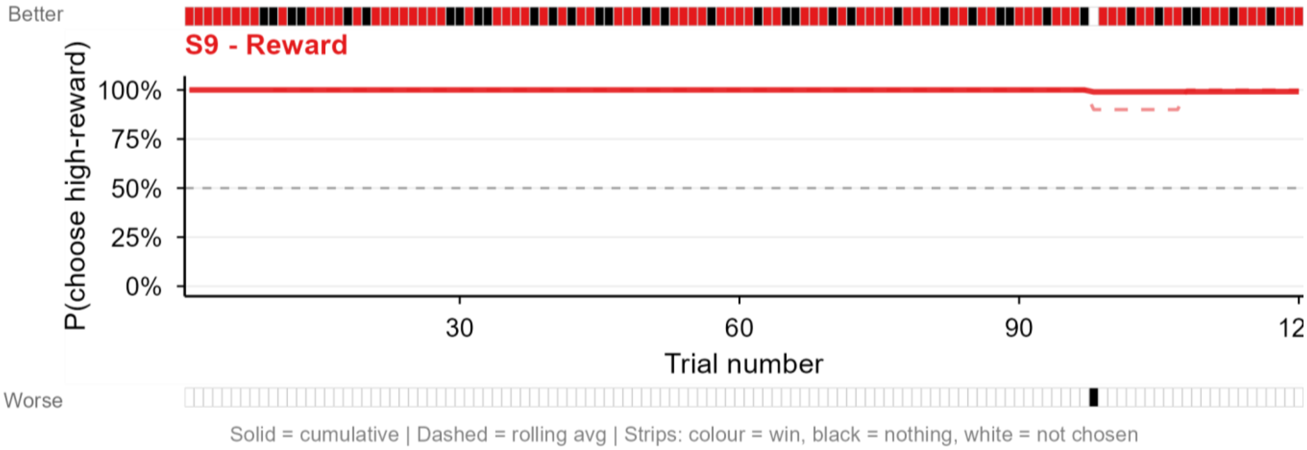

Better
S9 - Reward
P(choose high-reward)
100%
75%
50%
25%
0%
30
60
90
12
Trial number
Worse
Solid = cumulative | Dashed = rolling avg | Strips: colour = win, black = nothing, white = not chosen


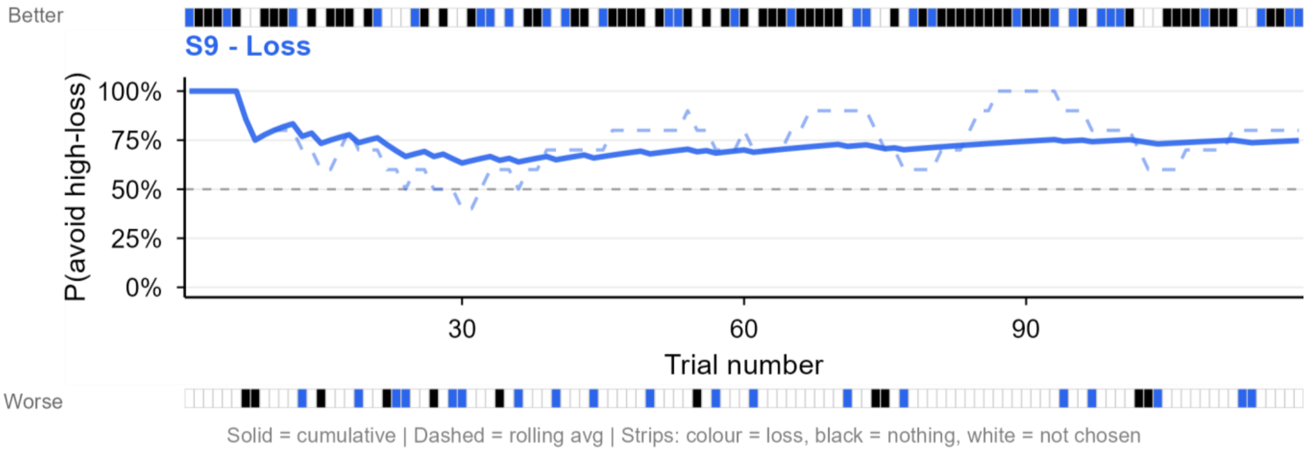

Better
S9 - Loss
P(avoid high-loss)
100%
75%
50%
25%
0%
30
60
90
Trial number
Worse
Solid = cumulative | Dashed = rolling avg | Strips: colour = loss, black = nothing, white = not chosen


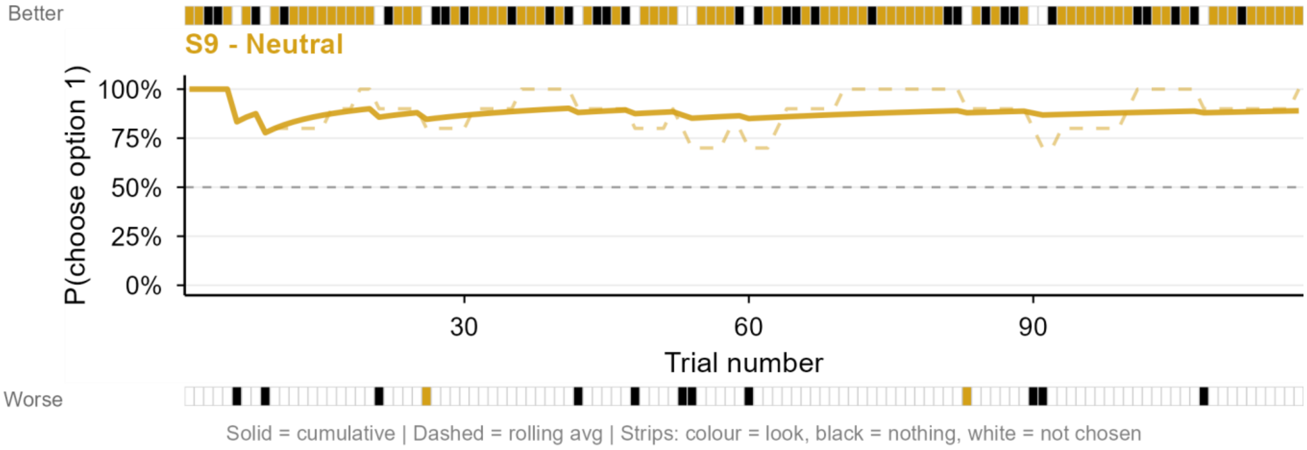

Better
S9 - Neutral
P(choose option 1)
100%
75%
50%
25%
0%
30
60
90
Trial number
Worse
Solid = cumulative | Dashed = rolling avg | Strips: colour = look, black = nothing, white = not chosen

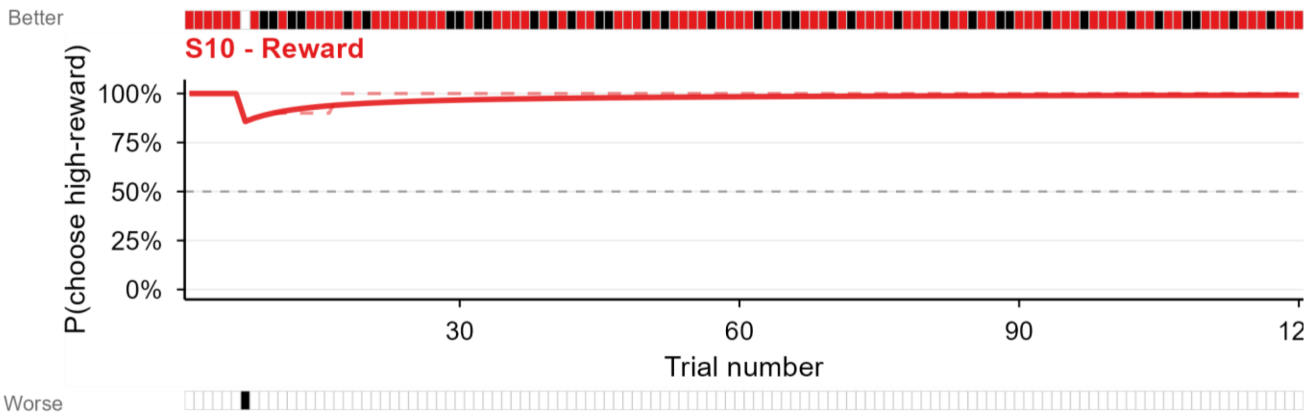


Solid = cumulative | Dashed = rolling avg | Strips: colour = win, black = nothing, white = not chosen

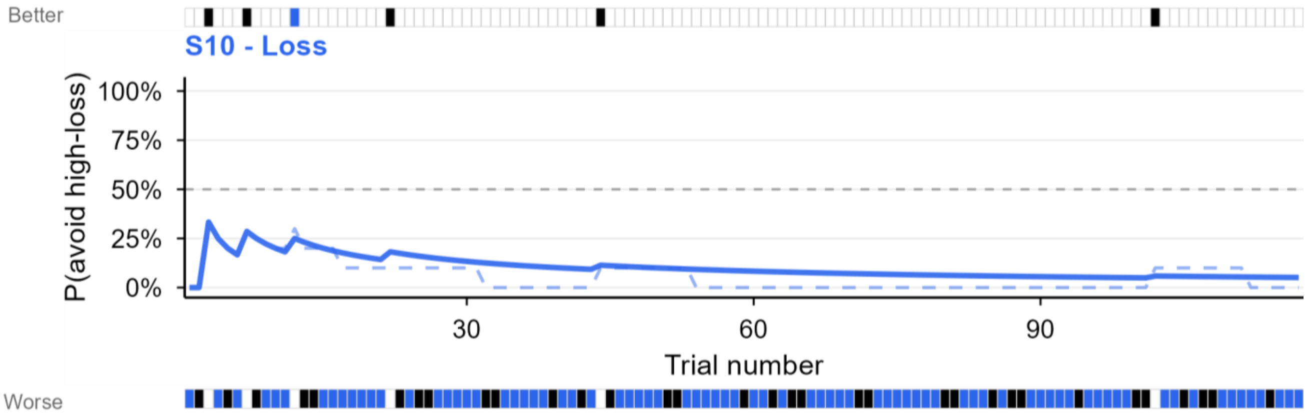


Solid = cumulative | Dashed = rolling avg | Strips: colour = loss, black = nothing, white = not chosen

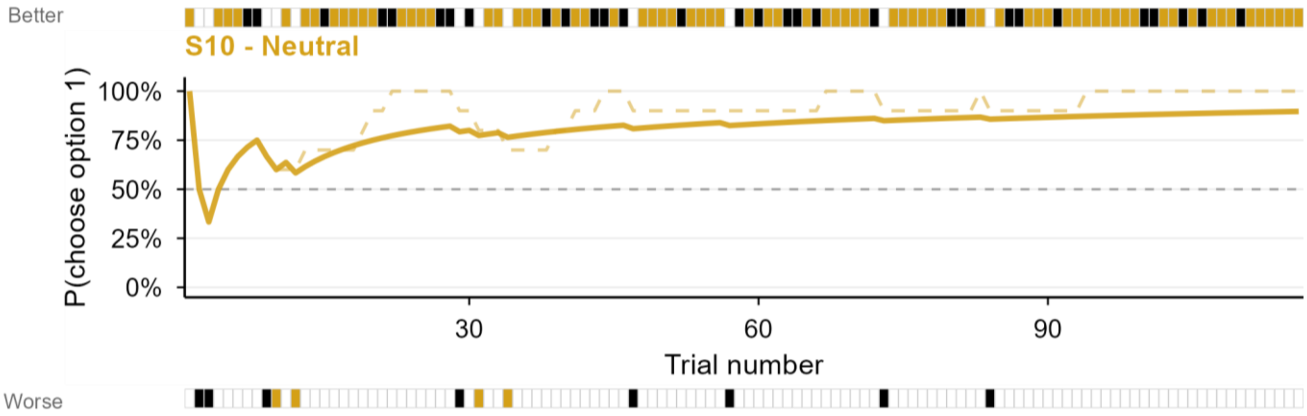


Solid = cumulative | Dashed = rolling avg | Strips: colour = look, black = nothing, white = not chosen

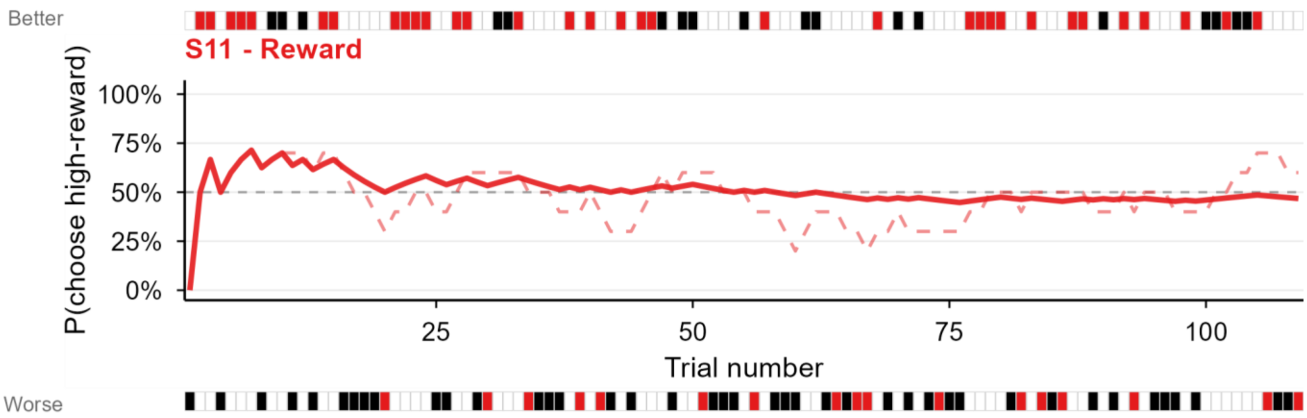


Solid = cumulative | Dashed = rolling avg | Strips: colour = win, black = nothing, white = not chosen

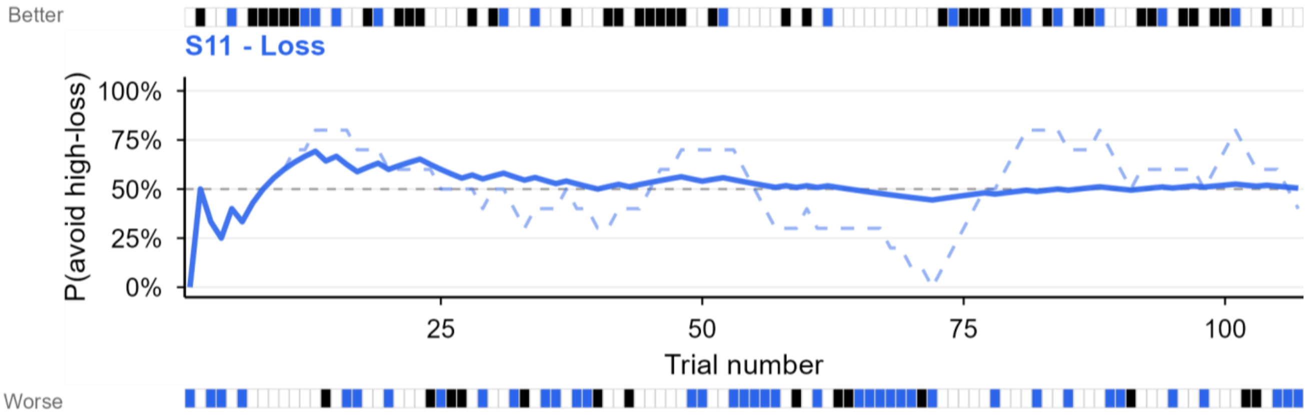


Solid = cumulative | Dashed = rolling avg | Strips: colour = loss, black = nothing, white = not chosen

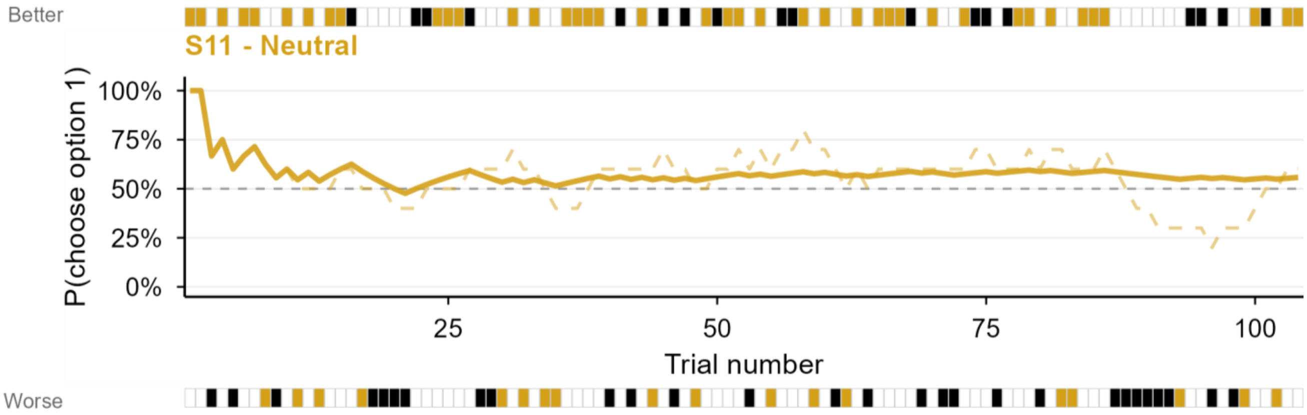


Solid = cumulative | Dashed = rolling avg | Strips: colour = look, black = nothing, white = not chosen

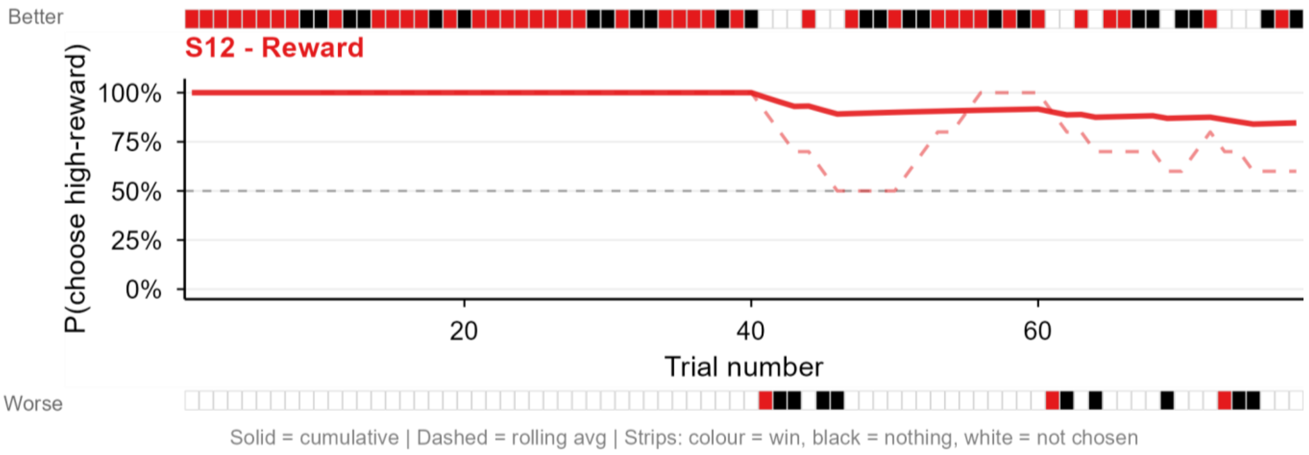
Better
S12 - Reward
P(choose high-reward)
100%
75%
50%
25%
0%
20
40
60
Trial number
Worse
Solid = cumulative | Dashed = rolling avg | Strips: colour = win, black = nothing, white = not chosen

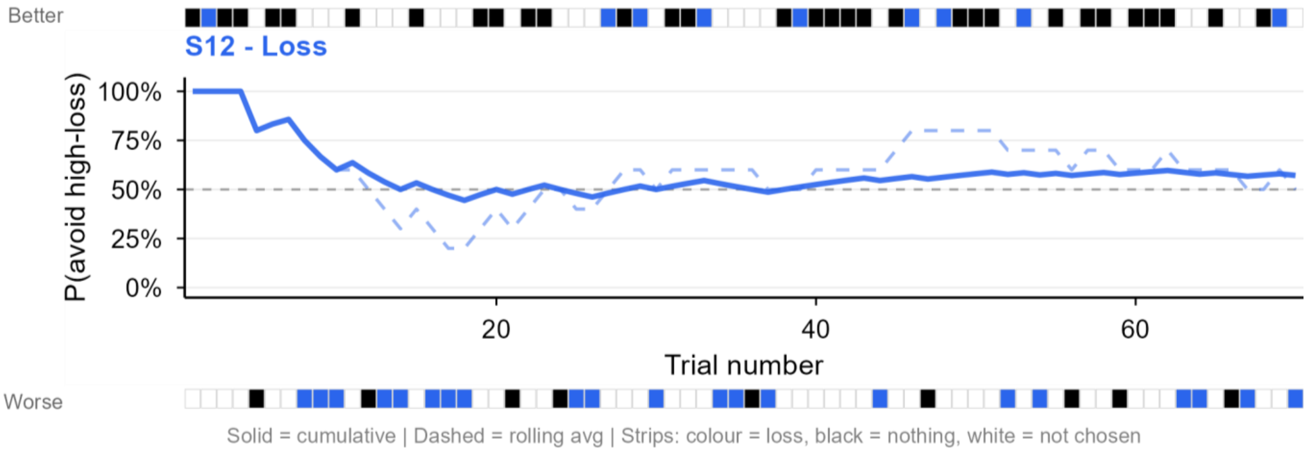
Better
S12 - Loss
P(avoid high-loss)
100%
75%
50%
25%
0%
20
40
60
Trial number
Worse
Solid = cumulative | Dashed = rolling avg | Strips: colour = loss, black = nothing, white = not chosen

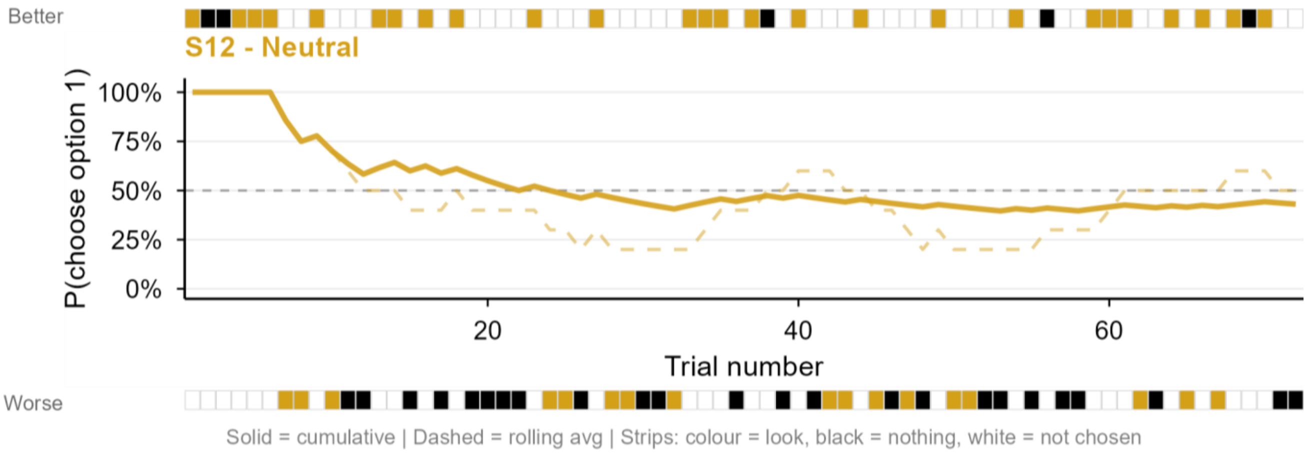
Better
S12 - Neutral
P(choose option 1)
100%
75%
50%
25%
0%
20
40
60
Trial number
Worse
Solid = cumulative | Dashed = rolling avg | Strips: colour = look, black = nothing, white = not chosen

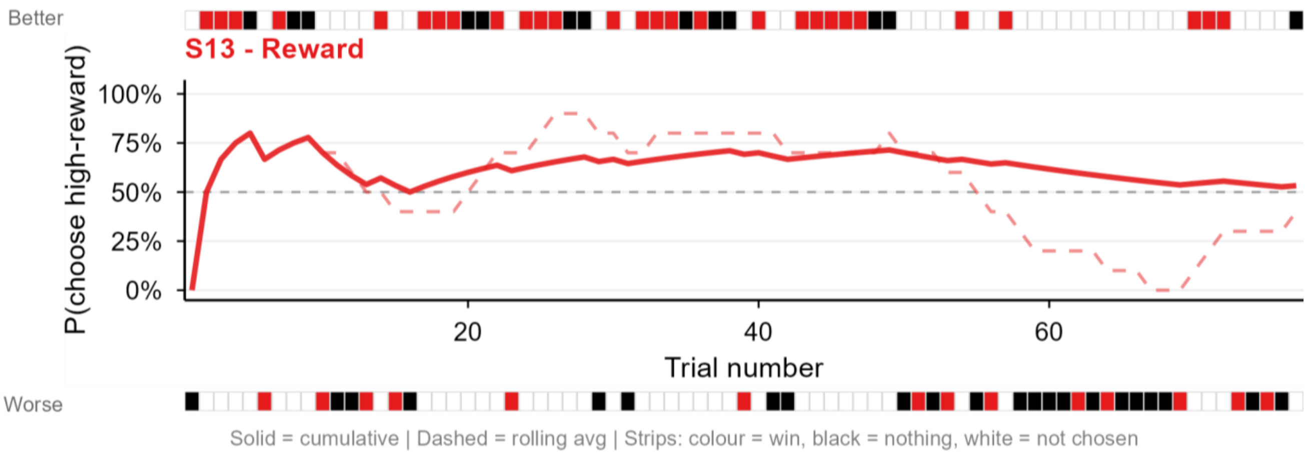
Better
S13 - Reward
P(choose high-reward)
100%
75%
50%
25%
0%
20
40
60
Trial number
Worse
Solid = cumulative | Dashed = rolling avg | Strips: colour = win, black = nothing, white = not chosen

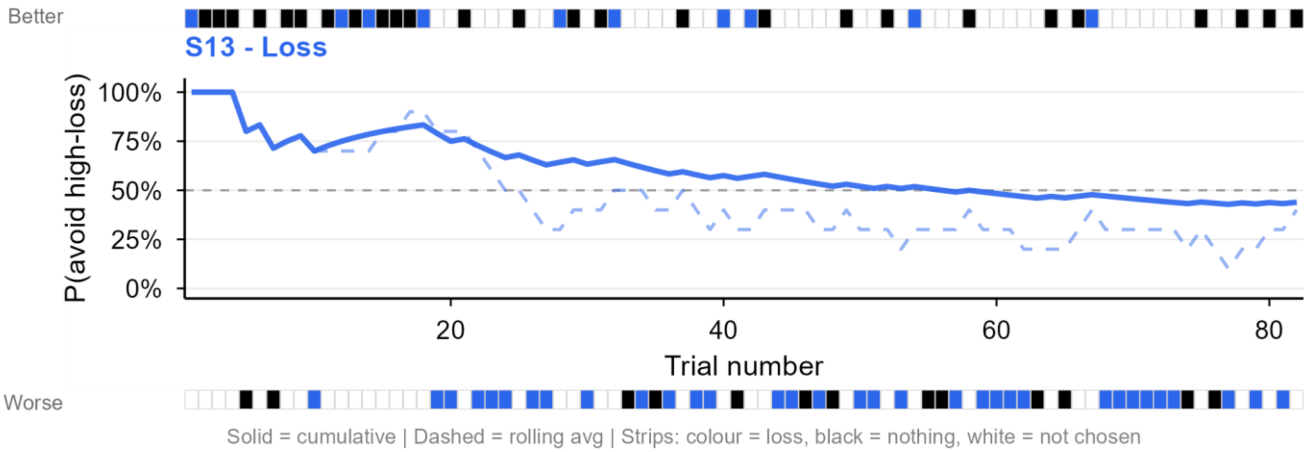
Better
S13 - Loss
P(avoid high-loss)
100%
75%
50%
25%
0%
20
40
60
80
Trial number
Worse
Solid = cumulative | Dashed = rolling avg | Strips: colour = loss, black = nothing, white = not chosen

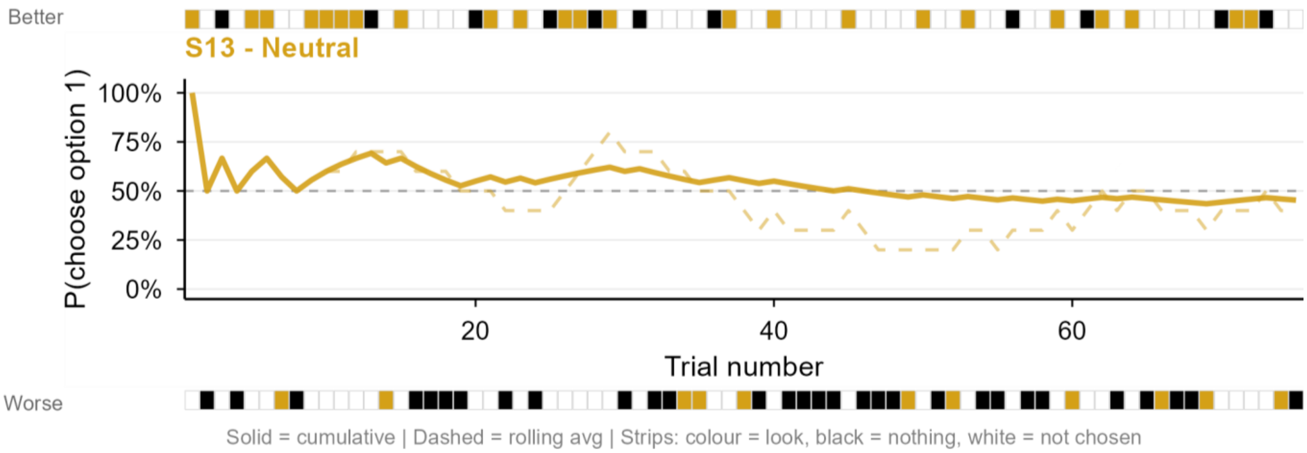
Better
S13 - Neutral
P(choose option 1)
100%
75%
50%
25%
0%
20
40
60
Trial number
Worse
Solid = cumulative | Dashed = rolling avg | Strips: colour = look, black = nothing, white = not chosen

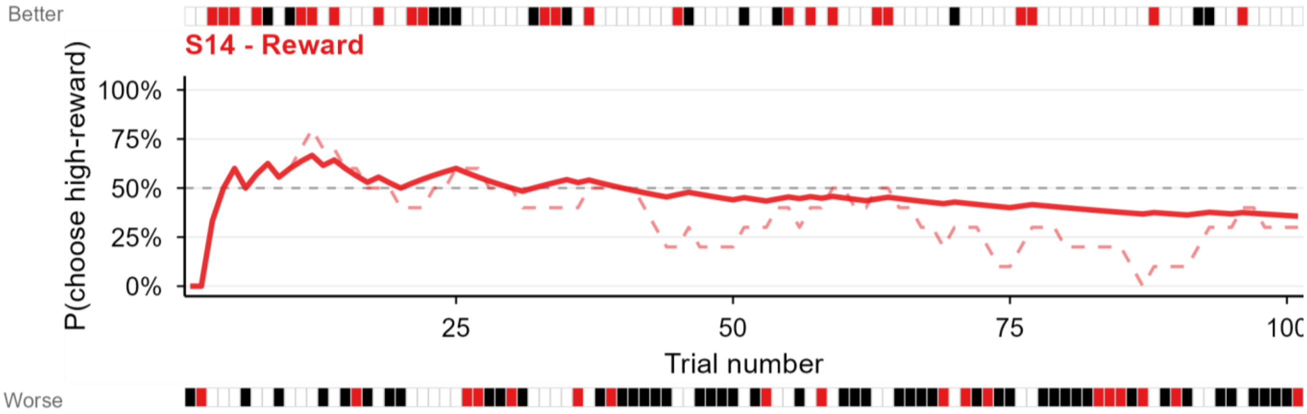


Solid = cumulative | Dashed = rolling avg | Strips: colour = win, black = nothing, white = not chosen

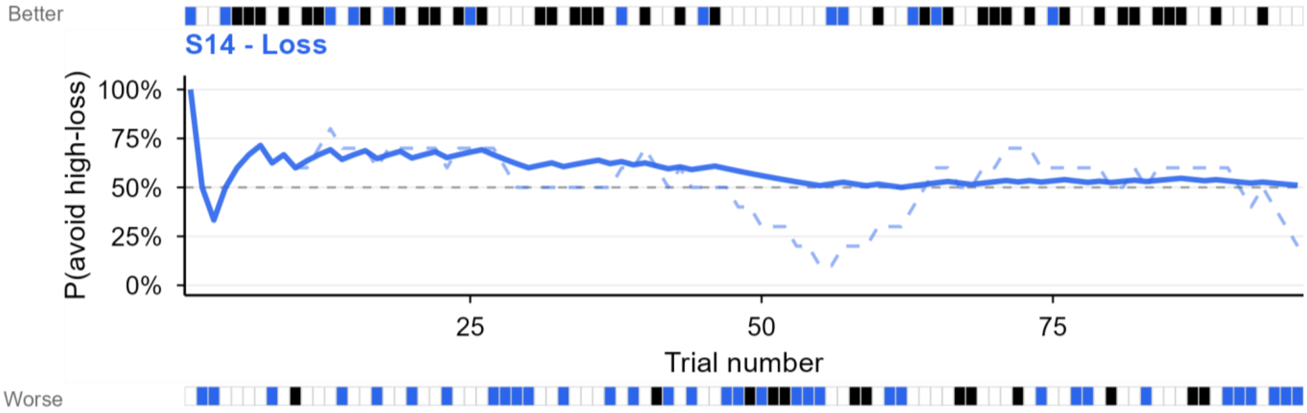


Solid = cumulative | Dashed = rolling avg | Strips: colour = loss, black = nothing, white = not chosen

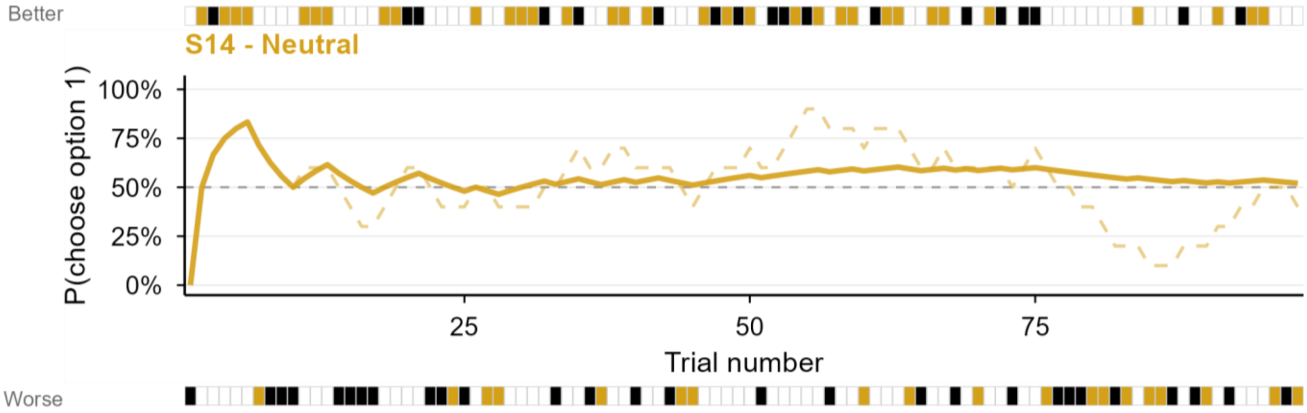


Solid = cumulative | Dashed = rolling avg | Strips: colour = look, black = nothing, white = not chosen

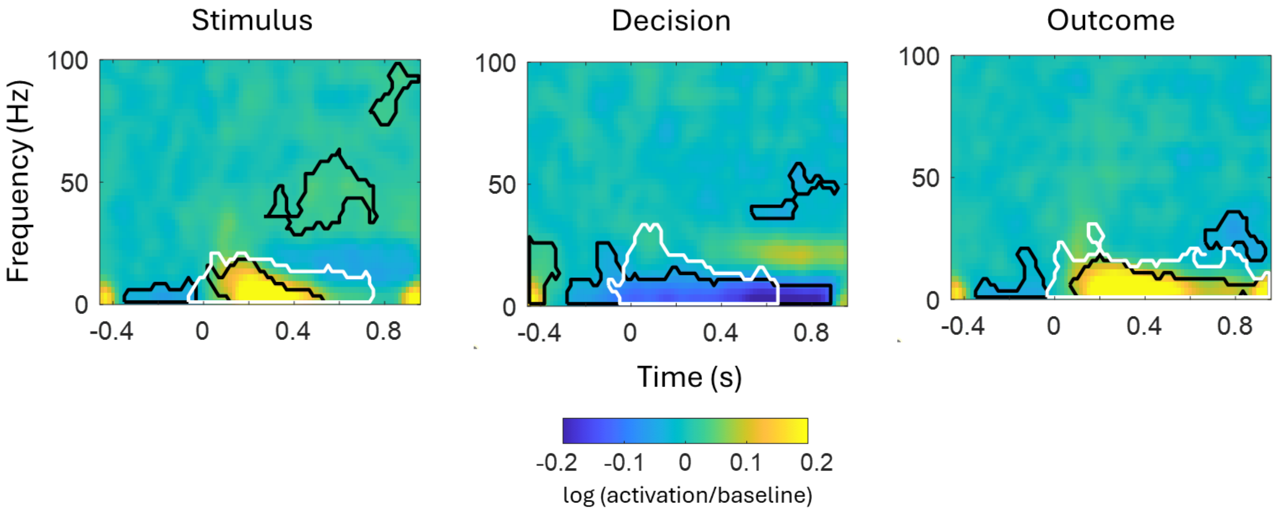


**Supplementary Figure S2. Induced responses to task events**

The black boundaries denote significant clusters of power modulations (FWE-corrected at the cluster level, p<0.05). The white boundaries denote significant clusters of inter-trial phase locking increase (FWE-corrected at the cluster level, p<0.05) marking areas where power modulations are likely to reflect evoked activity. The colour represents changes in log power compared to baseline.

**Supplementary Table S2. Post-hoc correlations between separate value regressors and VTA-LFP responses of Subject 3.**

| Time window (s) | Chosen option | High-value option | Low-value option |
|---|---|---|---|
| -0.5 to -0.43 | r=0.32, p=4.2e-04 | r=0.15, p=0.112 | r=-0.2, p=0.031 |
| -0.1 to 0.033 | r=-0.44, p=6.8e-07 | r=-0.29, p=0.001 | r= 0.24, p=0.01 |

The responses were averaged over significant clusters revealed by an F-test for all three regressors combined.

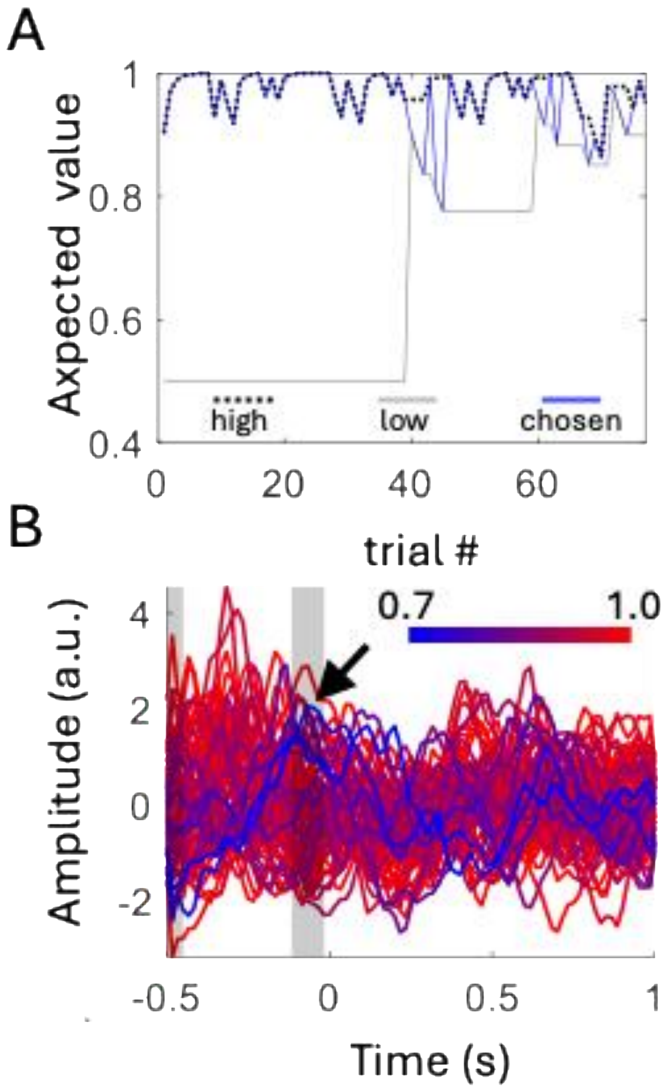


**Supplementary Figure S3. Value correlates in VTA-LFP responses of Subject 12 around the decision event (button press).** (**A**) Profile of the 'value of the chosen option' regressor across trials (blue line). The black dotted line shows the value of the high value option and the grey solid line – that of the low value option. The blue line alternates between the two depending on the choice. (**B**) Single-trial VTA-LFP responses, colour-coded by the chosen option's value (blue: lowest; red: highest). Shaded areas indicate significant clusters. The pattern of effect is similar to that shown for Subject 3 in Figure 6.
